# Assessing reliability of intra-tumor heterogeneity estimates from single sample whole exome sequencing data

**DOI:** 10.1101/440750

**Authors:** Judith Abécassis, Anne-Sophie Hamy, Cécile Laurent, Benjamin Sadacca, Hélène Bonsang-Kitzis, Fabien Reyal, Jean-Philippe Vert

## Abstract

Tumors are made of evolving and heterogeneous populations of cells which arise from successive appearance and expansion of subclonal populations, following acquisition of mutations conferring them a selective advantage. Those subclonal populations can be sensitive or resistant to different treatments, and provide information about tumor aetiology and future evolution. Hence, it is important to be able to assess the level of heterogeneity of tumors with high reliability for clinical applications.

In the past few years, a large number of methods have been proposed to estimate intra-tumor heterogeneity from whole exome sequencing (WES) data, but the accuracy and robustness of these methods on real data remains elusive. Here we systematically apply and compare 6 computational methods to estimate tumor heterogeneity on 1,697 WES samples from the cancer genome atlas (TCGA) covering 3 cancer types (breast invasive carcinoma, bladder urothelial carcinoma, and head and neck squamous cell carcinoma), and two distinct input mutation sets. We observe significant differences between the estimates produced by different methods, and identify several likely confounding factors in heterogeneity assessment for the different methods. We further show that the prognostic value of tumor heterogeneity for survival prediction is limited in those datasets, and find no evidence that it improves over prognosis based on other clinical variables.

In conclusion, heterogeneity inference from WES data on a single sample, and its use in cancer prognosis, should be considered with caution. Other approaches to assess intra-tumoral heterogeneity such as those based on multiple samples may be preferable for clinical applications.

## Introduction

Cancer is characterized by the presence of cells growing and dividing without proper control. In the 1970s, Nowell and colleagues suggested that tumor cells follow evolutionary principles, as any other biological population able to acquire heritable transformations [1]. This evolutionary framework has proven very useful in deepening our understanding of cancer aetiology [2].

A consequence of this progressive accumulation of mutations is intra-tumor heterogeneity. Indeed, when a new mutation occurs in a tumor cell and provides an evolutionary advantage, this cell tends to have a higher probability to survive and divide, hence seeding a new clonal population [3]. This new clone may supersede the whole tumor population, or coexist along it. This process results in a tumor made of a mosaic of clones. Next generation sequencing (NGS), in particular whole exome and whole genome sequencing (WES, WGS), can provide new insights into the heterogeneity and evolution of tumors. Indeed, early mutations shared among all cancer cells should be detected in more sequencing reads than mutations acquired later by only a fraction of the tumor cells. Thus it may be possible to estimate the intra-tumor heterogeneity (ITH) and reconstruct the clonal history of tumors from WES or WGS data, as reviewed by [3, 4, 5], and many computational methods have been developed for that purpose [6, 7, 8, 9]. We collectively refer to these methods as “ITH methods” in the following. Subclonal reconstruction from single cell sequencing has emerged as a new field, simplifying part of the inference problem, but raising other issues, related to technical limitations (high dropout rate) and high cost, possibly a limitation to the availability of large cohorts [10, 11, 12, 3].

Previous studies have reported that a large proportion of tumors are heterogeneous [13, 14, 15, 16], with various consequences for the patient. In particular, high ITH has been associated with treatment resistance and poor prognosis [17]. However, those results rely mostly on very detailed case studies involving only a small number of patients, with favorable experimental settings such as high coverage targeted sequencing on top of NGS, multiple sample collection (multi-site or longitudinal studies) [18, 19, 20] or even single-cell sequencing [21]. In the perspective of large-scale application in a clinical context, one needs to consider more accessible data with respect to cost and invasiveness for the patient, like moderate coverage WES on one sample per patient. A precise evaluation of existing ITH methods in this setting is needed to determine whether they allow us to find distinguishable patterns of heterogeneity and evolution of clinical relevance. Several large scale analyses have attempted to depict the evolutionary landscape of ITH in several cancer types [2], and to assess the prognostic power of ITH. In particular, using data from the cancer genome atlas (TCGA), a significant association between ITH and overall survival was found in at least one of the three studies [14, 13, 22] for 9 cancer types: breast invasive carcinoma (BRCA), kidney renal clear cell carcinoma (KIRC), brain lower grade glioma (LGG), prostate adenocarcinoma (PRAD), glioblastoma multiforme (GBM), head and neck squamous cell carcinoma (HNSC), ovarian serous cystadenocarcinoma (OV), uterine corpus endometrial carcinoma (UCEC), and colon adenocarcinoma (COAD). However, 5 of them were considered in another study with no significant result. In other cancer types, 2 studies consistently found no significant results for 3 cancer types: bladder urothelial carcinoma (BLAC), lung squamous cell carcinoma (LUSC) and stomach adenocarcinoma (STAD), and all 3 studies found no significant results for lung adenocarcinoma (LUAD) nor for skin cutaneous melanoma (SKM). A possible explanation for this discrepancy is that the studies base their analyses on different computational pipelines, from variant calling to ITH estimation, leading to different and sometimes contradictory results [22].

To clarify the robustness and consistency of different ITH methods, we perform a systematic benchmark of 18 computational pipelines for ITH estimates from a single WES sample per patient (combining 2 ways to call mutations, and 2 methods to assess copy number variations (only 3 out of 4 combinations were tested) with 6 ITH methods), using data from 1,697 patients with three types of cancer from the TCGA database (BRCA, BLCA, HNSC). We selected these cancer types following conclusions of Morris et al. [13], since HNSC, BRCA and BLCA are characterized by respectively high, intermediate and absence of prognostic power of ITH. We show that most existing ITH methods are very sensitive to the choice of mutations and copy number variations called, and that they can give very inconsistent results between each other. We highlight in particular that some methods are influenced by confounding factors such as tumor purity or mutation load. Finally, we show that although ITH measured by some computational pipelines have a weak prognostic power on some cancer types, the prognosis signal is not robust across methods and cancer types, and is confounded with informations available in standard clinical data. To further characterize those inconsistencies, we report results for ITH methods on 7 WES samples associated with single cell sequencing allowing to have an estimate of the ground truth. As a conclusion, we suggest that results of ITH analysis from single sample WES data with current computational pipelines should be manipulated with caution, and that more robust methods or protocols are likely to be needed for clinical applications.

## Materials and methods

### Data

We downloaded data from the GDC data portal https://portal.gdc.cancer.gov/ for 3 cancer types (BLCA - 351 patients, BRCA - 904 patients, HNSC - 442 patients). We gathered annotated somatic mutations, both raw variant calling output, whose access is restricted and public mutations, from the new unified TCGA pipeline https://docs.gdc.cancer.gov/Data/Bioinformatics_Pipelines/DNA_Seq_Variant_Calling_Pipeline/, with alignment to the GRCh38 assembly, and variant calling using 4 variant callers: MuSe, Mutect2, VarScan2 and SomaticSniper. Instructions for download can be found in the companion Github repository (https://github.com/judithabk6/ITH_TCGA). RNAseq data used to compute immune signatures were downloaded through TCGABiolinks [23], and we downloaded clinical data from the CBIO portal [24].

### Copy number calling and purity estimation

We obtained copy number alterations (CNA) data from the ASCAT complete results on TCGA data partly reported on the COSMIC database [25, 26]. We then converted ASCAT results on hg19 to GRCh38 coordinates using the segment liftover Python package [27]. ASCAT results also provide an estimate of purity, which we used as input to ITH methods when possible. Other purity measures are available [28]; however we selected the ASCAT estimate to ensure consistency with CNV data.

The calls of allele-specific copy number and purity from ABSOLUTE [29] were downloaded from the GDC data portal https://gdc.cancer.gov/about-data/publications/pancanatlas on August 18th 2019. They were converted to GRCh38 as the ones from ASCAT.

### Variant calling filtering

Variant calling is known to be a challenging problem. It is common practice to filter variant callers output, as ITH methods are deemed to be highly sensitive to false positive single nucleotide variants (SNVs). We filtered out indels from the public dataset, and considered the union of the 4 variant callers output SNVs. For the protected data, we also removed indels, and then filtered SNVs on the FILTER columns output by the variant caller (”PASS” only VarScan2, SomaticSniper, ”PASS” or ”panel of normals” for Mutect2, and ”Tier1” to ”Tier5” for MuSe). In addition, for all variant callers, we removed SNVs with a frequency in 1000 genomes or Exac greater than 0.01, except if the SNV was reported in COSMIC. A coverage filter was added, and we kept SNVs with at least 6 reads at the position in the normal sample, of which 1 maximum reports the alternative nucleotide (or with a variant allele frequency (VAF) *<*0.01), and for the tumor sample, at least 8 reads covering the position, of which at least 3 reporting the variant, or a VAF*>*0.2. The relative amount of excluded SNVs from protected to public SNV sets varied significantly between the 3 cancer types (see Table S1). All annotations are the ones downloaded from the TCGA, using VEP v84, and GENCODE v.22, sift v.5.2.2, ESP v.20141103, polyphen v.2.2.2, dbSNP v.146, Ensembl genebuild v.2014-07, Ensembl regbuild v.13.0, HGMD public v.20154, ClinVar v.201601. We further denote the filtered raw mutation set as ”Protected SNVs” and the other one, which is publicly available, as ”Public SNVs”

**Table 1:** Main characteristics of ITH methods tested. The mean runtime is the mean time to process a TCGA sample. The success rate is the fraction of TCGA samples for which the method produced an output without error, with ASCAT calls as input only. The MATH score was computed in one step for all samples, using a table containing all mutations for all samples; the operation lasted 3.21s (std. 47.6 ms) for the protected dataset, and 3.39s (std. 11ms) for the public dataset. All time measurements were measured on a single cluster node with a 2.2 GHz processor and 3GB of RAM.

| Method | CNA as input | Purity as input | Outputs tree(s) | Reference | Mean (std) runtime protected in seconds | Mean (std) runtime public in seconds | Success rate (protected) | Success rate (public) |
| --- | --- | --- | --- | --- | --- | --- | --- | --- |
| MATH | no | no | no | [30] | << 1 | << 1 | 100% | 100% |
| EXPANDS | yes | no | no | [9] | 891 (604) | 267 (258) | 89% | **71%** |
| PyClone | yes | yes | no | [6] | 7,035 (8,464) | 1,414 (1,415) | 95% | 99% |
| SciClone | yes | no | no | [7] | 62 (48) | 41 (51) | 92% | **78%** |
| PhyloWGS | yes | yes | yes | [8] | 13,258 (9,058) | 4,730 (4,139) | 95% | 97% |

### ITH methods

#### Published methods

We consider four published ITH methods: SciClone [7], PhyloWGS [8], PyClone [6] and EXPANDS [9]. In addition, we consider the MATH score [30] as a simple indicator of ITH, as well as a baseline ITH method described below. All computations were stopped after running 15 hours. This threshold was chosen to get results for most samples (*>* 95% when time was the limiting factor) for most methods while saving computational resources. Mean and standard deviation (std) of runtimes were computed for each method with each input mutation set separately. All parameters used for each method are detailed in the companion public Github repository containing all the commands https://github.com/judithabk6/ITH_TCGA. To ensure comparison, the runtimes were only performed on runs with ASCAT copy number calls.

We performed post-treatment to keep only clones with at least 5 SNVs, except for samples in which all clones were under 5 SNVs when all clones were considered. After running each ITH method we extracted 5 features to characterize ITH in a sample: the number of clones, the proportion of SNVs that belong the the major clone, the minimal cellular prevalence of a subclone, the Shannon index of the clonal distribution, and the cellular prevalence of the largest clone in terms of number of SNVs.

#### Consensus (CSR)

We computed a consensus of several ITH methods using the open source package CSR available at https://github.com/kaixiany/CSR. This method relies on matrix factorization to output a consensus clustering. We computed two separate consensus (for protected and public data), using as input the results of PyClone, SciClone, PhyloWGS, EXPANDS and baseline. MATH estimates were not well suited for the consensus. For each run, we ran matrix factorization for a maximum of 500 seconds.

### Clinical variables

For each cancer type, we collected clinical variables from the CBIO Portal according to the following conditions: (i) categorical variables were one-hot encoded, and each level was kept if it involved at least 50 patients, and at most 50 patients had another level of the same variable; (ii) we kept numerical variables available for every patient; and (iii) in addition, we only kept the variables (if numerical) or the levels (categorical) which were significantly associated with overall survival by a single-variable cox model estimated with the Python package lifelines [31] after Benjamini-Hochberg correction for multiple hypothesis testing [32]. Tables S2, S3, and S4 summarize the clinical variables retained for each cancer type.

**Table 2:**
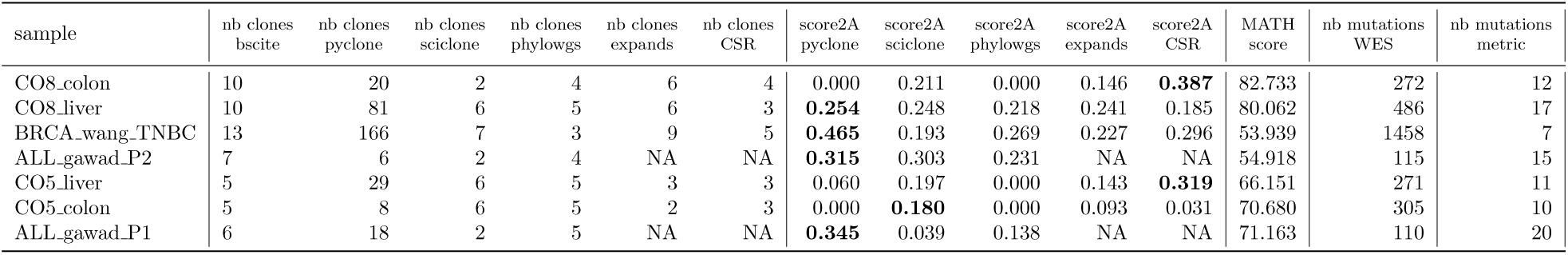
Results on the single cell-WES dataset.

### Survival regression

#### Model

To estimate the prognosis power of a set of features, we use a survival SVM model [33]. Survival SVM maximizes a concave relaxation of the concordance between the predicted survival ranks and the original observed survival, regularized by a Euclidean norm penalty. Formally, given a training set of *n* patients with survival information (**x***_i_, y_i_, δ_i_*)*_i_*_=1_*_,…,n_*, where **x***_i_ ∈* ℝ*^p^* is a vector of *p* features for patient *i*, *y_i_ ∈* ℝ is the time, and *δ_i_ ∈* {0, 1} indicates the event (*δ_i_* = 1) or censoring (*δ_i_* = 0), a survival SVM learns a linear score of the form *f* (**x**) = **w***^T^***x** for any new patient represented by features **x** *∈* ℝ*^p^* by solving:

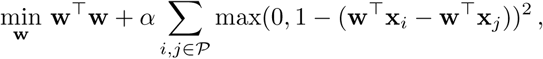

where *P* = {(*i, j*) ∈ [1, n]^2^ | *y_i_ ≥ y_j_ ^ δ_j_* = 1} is the set of pairs of patients (*i, j*) which are comparable, that is, for which we are certain that patient *i* lived longer than patient *j*. Intuitively, the loss penalizes the cases where patient *i* survives longer than patient *j* but the opposite is predicted by the model. For all computations, we used the function FastSurvivalSVM in the Python Package scikit-survival [34], with default parameters. The model was trained and tested using a 5-fold cross-validation procedure.

#### Evaluation procedure

To assess the accuracy of a survival regression model, we use the concordance index (CI) between the predicted score and the true survival information on a cohort with survival information. Given such a cohort (**x***_i_, y_i_, δ_i_*)*_i_*_=1_*_,…,n_*, the CI measures how concordant the predicted survival times *s_i_* = *f* (**x***_i_*) are with the observed survival times *y_i_* for comparable pairs of patients:

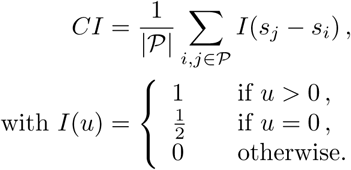

In practice, we compute an approximation of CI with the function concordance.index from the R package survcomp [35], using the noether method [36], and the associated one-sided test to compare CI to 0.5, which is the mean CI obtained with a random predictor. To compare CI’s of different methods, we use a paired Student t-test for dependent samples implemented in the function cindex.comp from the same package. In both test settings, we aggregate p-values from each of the five cross-validation folds using the Fisher method from Python package statsmodels, and apply a Benjamini-Hochberg correction [32] to correct for multiple testing.

### Immune signatures

We normalized RNAseq raw count data using a variance stabilizing transformation (VST) implemented in the Deseq2 R package [37], treating each cancer type separately. We mapped genes from Bindea et al. [38] to Ensembl GeneIds present in the TCGA matrix using EntrezId match table downloaded from Biomart [39] on March 26th 2018. Out of 681 EntrezId (577 unique), 31 (24 unique) were not matched to an Ensembl Id with associated gene expression in the TCGA RNAseq data. Each signature was then computed by averaging the VST output value for the relevant Ensembl Id for each TCGA sample. The resulting signatures we used can be found as Supplementary Table S5. For analysis purposes, we use the complementary to the maximal value in the cohort so that the content in immune cells varies in the same direction as tumor purity and remains a positive quantity. We denote those new variables with the prefix inv, e.g., for patient *i* in the BRCA cohort we define

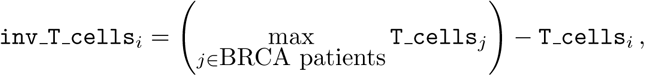

where T_cells*_i_* represents the signature for T cells estimated as explained above.

### Correlations

We assessed correlations using Pearson’s correlation coefficient. We computed the associated significance (for the null hypothesis that the correlation coefficient is 0) using the scipy.stats.pearsonr function, and we corrected the significance for multiple testing using the Benjamini Hochberg procedure at *FDR ≤* 0.05.

#### Comparison metrics

In addition to the correlations of the number of clones between methods, we have implemented three metrics derived from [40] to compare ITH methods together:

**Score1B** measures the adequacy between one number of clones *J*_1_ and another number of clones *J*_2_. It is computed as 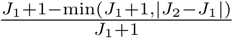.

**Score1C** is the Wasserstein distance between two clusterings, defined by the CCFs of the different clones and their associated weights (proportion of mutations), implemented as the function stats.wasserstein distance in the Python package scipy.

**Score2A** measures the correlation between two binary co-clustering matrices in a vector form, *M*_1_ and *M*_2_. It is the average of 3 correlation coefficients:

**Pearson correlation coefficient** 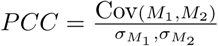, implemented as the function pearsonr in the Python package scipy,

**Matthews correlation coefficient** 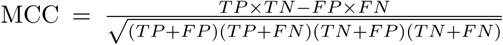, implemented as the function metrics.matthews corrcoef in the Python package scikit-learn,

**V-measure** is the harmonic mean of a homogeneity score that quantifies the fact that each cluster contains only members of a single class, and a completeness score measuring if all members of a given class are assigned to the same cluster [41]; here the classes are the true clustering. We used the function v measure score in the Python package scikit-learn.

Before averaging, all those scores were rescaled between 0 and 1 using the score of the minimal score between two ”bad scenarios”: all mutations are in the same cluster, or all mutations are in their own cluster (*M_pred_* = **1***_N×N_* or *M_pred_* = 𝕀*_N×N_*).

All scores as asymmetrical and were hence computed twice. In the case of score2A, only the mutations present in the two reconstructions were considered.

### WES and single cell paired dataset

#### Data availability and preprocessing

The raw data for 7 normal-tumor WES samples analyzed jointly with matching single cell sequencing [42] were downloaded from the NCBI SRA platform https://www.ncbi.nlm.nih.gov/sra and processed into fastq format using the tool fastq-dump for the two acute lymphoblastic leukemia (ALL) patients (accession numbers: SRR1517761, SRR1517762, SRR1517763, SRR1517764) [43], or directly downloaded in the fastq format from the EBI ENA platform https://www.ebi.ac.uk/ena for the Triple Negative Breast Cancer patient (TNBC) [44] (accession number: SRR1163508 and SRR1298936), and the two samples (primary tumor and liver metastasis) from the two colorectal cancer patients (CRC) [45] (accession number: SRR3472566, SRR3472567, SRR3472569, SRR3472571, SRR3472796, SRR3472798, SRR3472799, SRR3472800).

All normal-tumor pairs underwent a pipeline of analysis including alignment with BWA-MEM [46] with options ”-k 19 -T 30 -M”, filtering of reads based on target intersection, mapping quality and PCR duplicates removal, using Picard [47], Bedtools [48] and Samtools [49], and preprocess using GATK [50] for local realignment around indels, and base score recalibration. Variant calling was performed using Mutect2 [51], and variants filtered under the same rules as used for the TCGA (only ”PASS” variants, and minimal covering rules), and copy number assessed with Facets [52]. SNVs used in the analysis with B-SCITE [42], passing the covering filters but not recovered by this pipeline were added to the final variant list. Those variants and the copy number profile were then passed to PyClone, SciClone, PhyloWGS and Expands for ITH deconvolution.

#### Evaluation metrics

To measure the accuracy of subclonal reconstructions from the WES data only using different methods, we compared these reconstructions to the reconstruction obtained by B-SCITE using both WES and single cell sequencing [42]. To quantify the similarity of the different reconstruction results, we compared the number of clones, and for the common mutations, the metric 2A, used in [40] and redefined above.

## Results

### Assessing ITH on TCGA samples

We collected somatic mutation information from 1,697 TCGA patients with BLCA (*n* = 351), BRCA (*n* = 904), and HNSC (*n* = 442). We selected these three cancer types following conclusions of Morris et al. [13], since HNSC, BRCA and BLCA are characterized by respectively high (hazard ratio, HR=3.75, p=0.007 in multivariate Cox model), intermediate (HR=2.5, p=0.15) and absence (HR=1.05, p=0.91) of prognostic power of ITH. For each patient, we collected two sets of mutations based respectively on protected and public SNV sets. The protected set corresponds to raw variant calling outputs, with an extra filtering step described in Methods. The public set corresponds to publicly available SNV calls, filtered from the raw variant calling outputs to only retain somatic mutations with very high confidence, in order to ensure patients’ anonymity. Supplementary Table S1 summarizes some statistics on the number of mutations per sample for each cancer type.

We assess ITH in each sample using 6 representative computational methods: PyClone [6], SciClone [7], PhyloWGS [8] EXPANDS [9], the mutant-allele tumor heterogeneity (MATH) score [30], and CSR [16], a method providing a consensus of all of the above results (except MATH which is not compatible, see Methods). Table 1 summarizes some important properties of the different methods, which might be helpful for designing future studies and selecting the appropriate tool. All methods but MATH take as input the CNA information in addition to a set of somatic mutation VAFs. PyClone and PhyloWGS also take purity as input. All input has to be pre-computed by third-party approaches. While MATH is a single quantitative measure of ITH based on differences in the mutant-allele fractions among mutated loci, all 6 other methods produce more details such as the number of subclones and their respective proportions in the tumor. In particular, PhyloWGS outputs a lineage tree connecting the subclones.

We tested each method three times: on each sample for the two mutation sets combined with ASCAT calls for purity and copy number, and combined with ABSOLUTE calls for the protected mutation set. We observed that some methods failed to produce an output on some samples, for different reasons (see success rate for each method in Table 1). EXPANDS produces an error for 30% of the samples, mostly for tumors with high purity or very few CNAs. SciClone fails to provide an output for samples with an insufficient number of SNV in regions without CNA or LOH event. PyClone and PhyloWGS non completion cases were caused by a too long runtime.

As shown in Figure 1, there is little overlap between the samples where each method fails. Out of 1,697 initial TCGA samples, all methods produced an output for the three runs on only 686 samples (296 BRCA, 178 BLCA, 212 HNSC). Those failure cases unveil indications of each method’s limitations, in particular EXPANDS and SciClone. In the following we restrict our analysis to those 686 samples. One can note that there is more difference between public and protected results for BRCA samples; this is expected as the number of mutations in those two sets is more different for this cancer type, as shown in S1.

**Figure 1:**
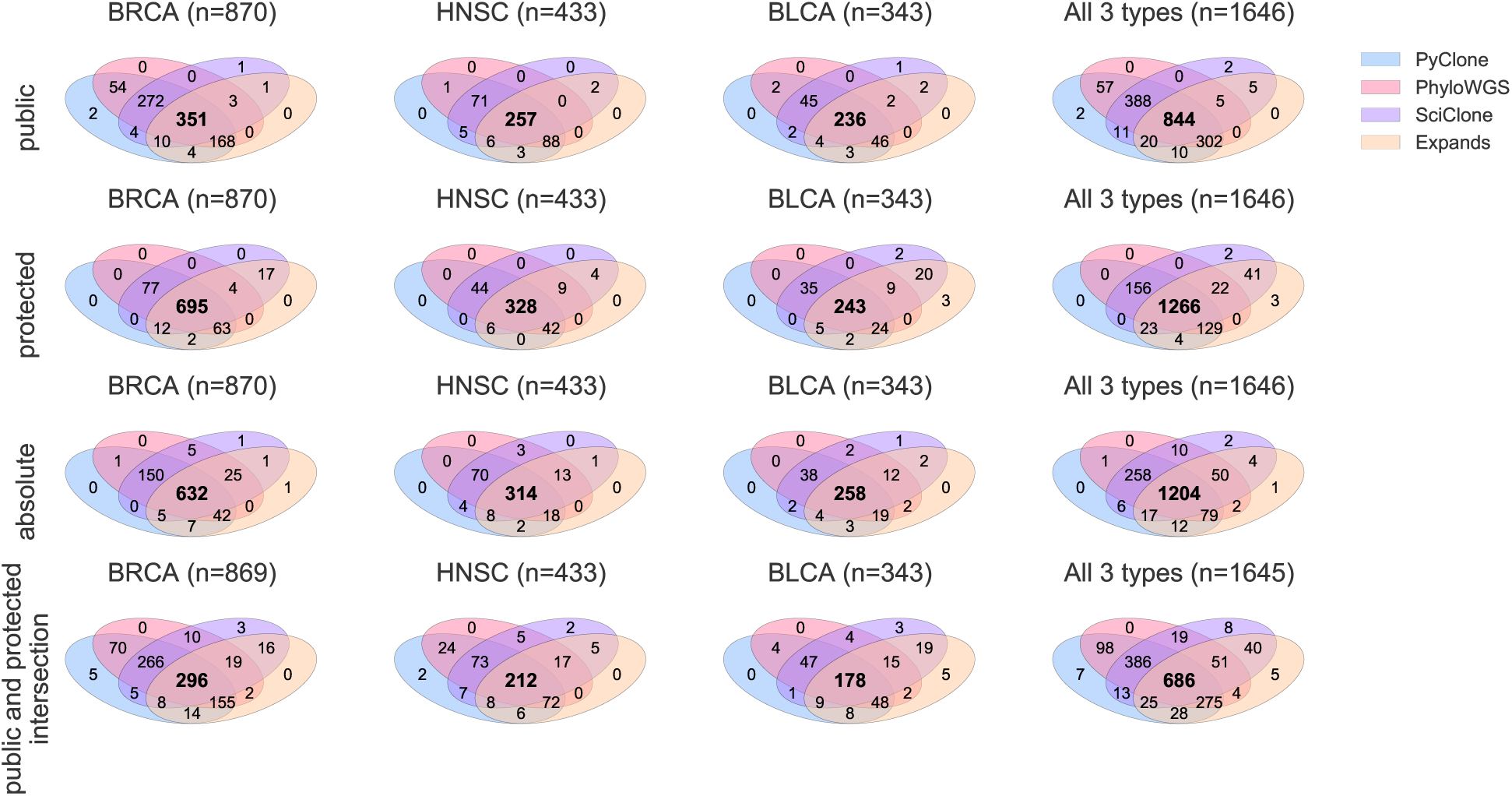
Intersection of successful runs among the 4 considered ITH methods. The upper venn diagrams concern runs with the public input SNV set, the second line with the protected, and the third the overall intersection, as results with both sets are necessary for a proper and rigorous comparison.

In addition to failures, we observed that the runtime varies significantly between methods (Table 1). As shown on Figure 2, the run time of different ITH methods increases with the number of somatic mutations. PyClone and PhyloWGS runtime rises very quickly with the number of mutations in tumor sample, which can be a limitation for applications to heavily mutated tumors.

**Figure 2:**
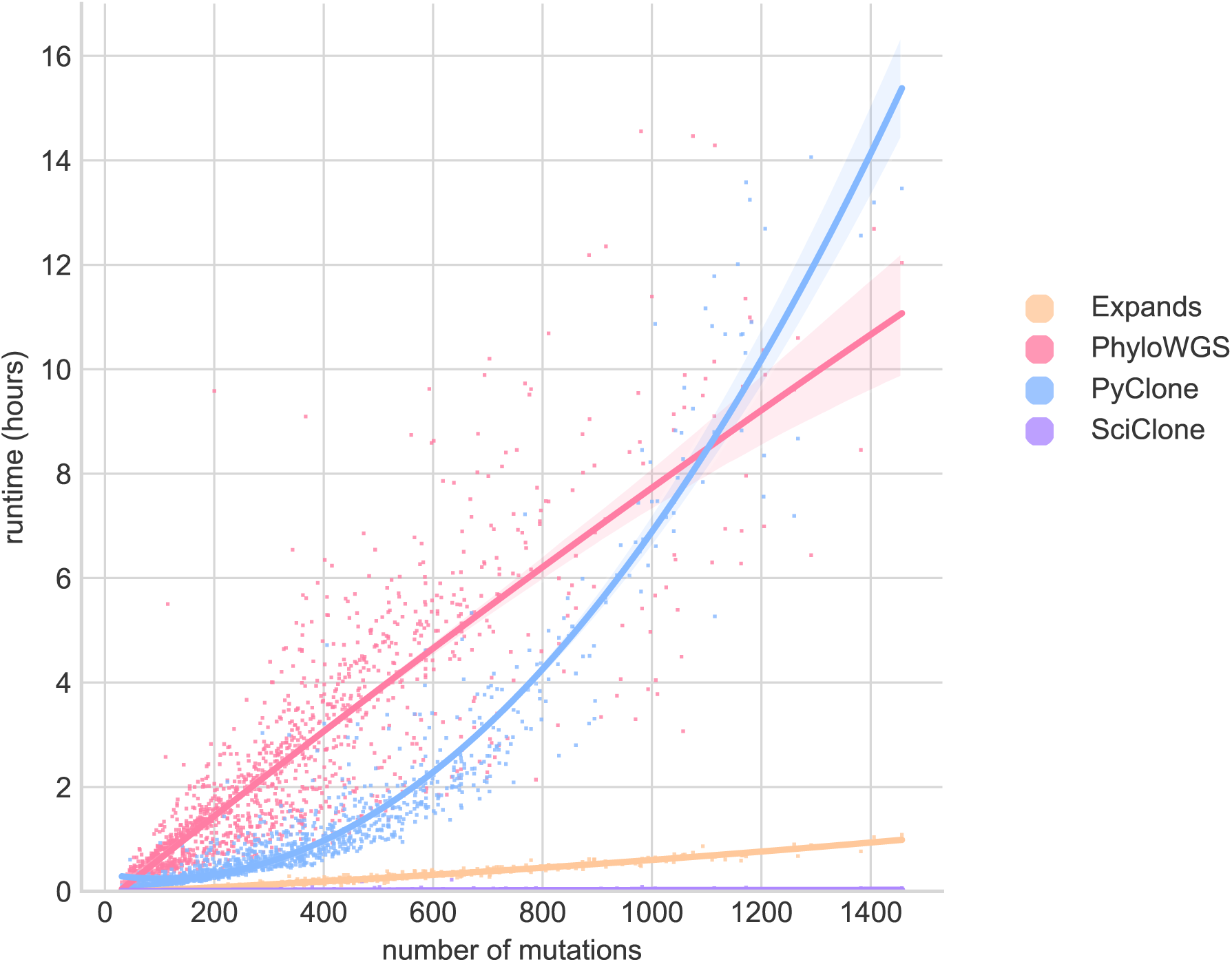
Runtime of the different ITH methods as a function of the number of mutations in each sample. Lines represent second degree polynomial fit with shaded regions are 95% confidence-intervals

### Methods quantifying ITH exhibit inconsistent results

As a first evaluation of ITH methods in the absence of ground truth, we assess the agreement between methods, with a focus on the number of clones. Each method except MATH outputs an estimated number *S* of subclonal populations, ranging from *S* = 1 for an homogeneous, clonal tumor to any positive number for an heterogeneous one. Figure 3 presents the distribution of estimated clonality among all samples for each approach and each SNV set, and each copy number calling method. We observe large differences between methods, as well as between SNV sets: for instance, over all samples, the percentage of estimated clonal tumors (*S* = 1) varies from 4% (for PhyloWGS on protected data) to 57 % (for PyClone on public data). Moreover, the number of estimated populations can vary strikingly with the mutation set used, but not really with the different input copy number. There is a clear trend among all methods to yield higher ITH estimates with the protected mutation set. PhyloWGS and EXPANDS (and CSR) are the only methods that detect ITH in almost all tested samples with the protected mutation set.

**Figure 3:**
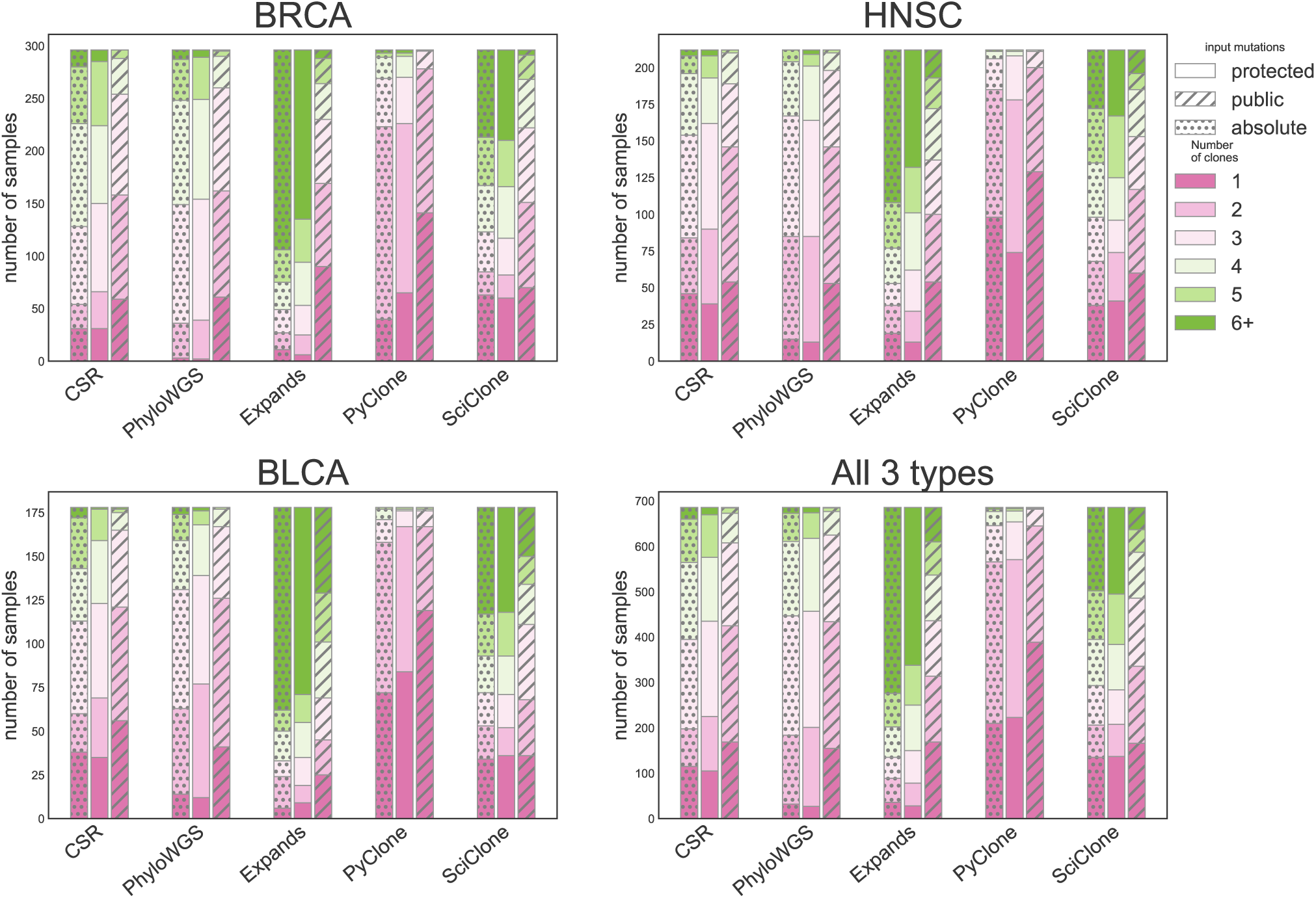
Distribution of number of clones called by ITH methods with public and protected mutations sets as inputs. Distribution of the number of subclones for the tested ITH methods, and 2 alternative input mutation sets for samples in the different cancer types and 2 different copy number methods for the protected mutation set. MATH could not be included in this analysis as this method does not estimate a number of clones.

Another way to compare methods is to consider correlations (Pearson’s r) between the estimated numbers of populations. This allows us to include the MATH score in the evaluation, considering it as an increasing function of heterogeneity just like the number of populations. In addition, we add to the comparison 5 measures directly extracted from the NGS analysis, namely, the number of mutations in the protected and in the public sets, the percentage of non-diploid cells (estimated by ASCAT and ABSOLUTE), the purity (estimated by ASCAT and ABSOLUTE), and the *inv T cell* (estimated from gene expression signatures). Results are presented in Figure 4.

**Figure 4:**
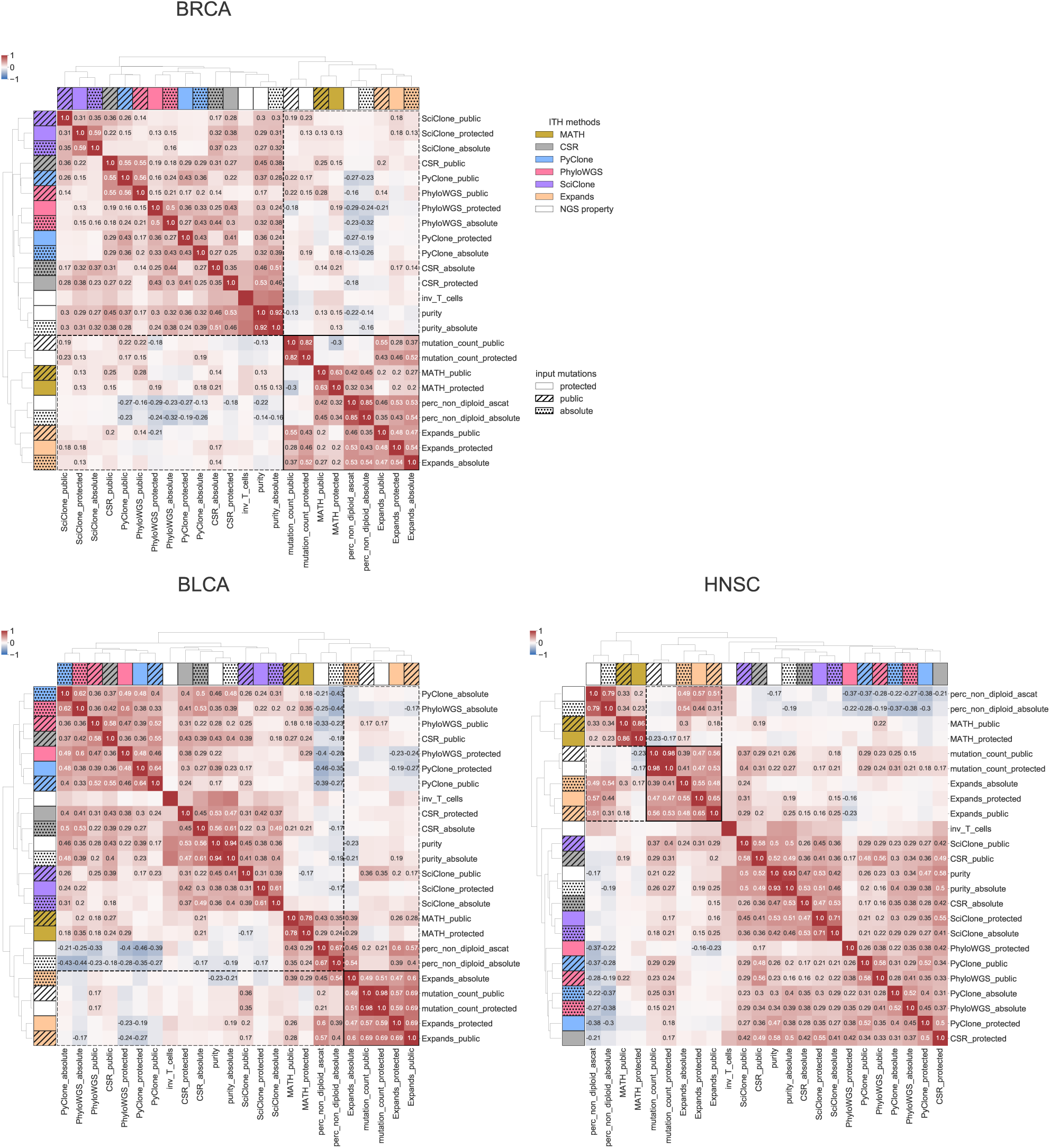
Correlation between various measures of ITH (MATH score, and number of subclones for the other methods), and other potential confounding variables measured using WES and trancriptomics data. Row and color label represent the method used, with white for the genomic measures not involving ITH. Hatches correspond to public mutation sets. Heatmap colors represent the value of the Pearson’s *r*, which is written numerically whenever it is significantly different from 0 (*FDR* 0.05 after Benjamini Hochberg correction for multiple tests). We can observe clustering tendencies stable across the 3 cancer types. One of them is highlighted in black lines.

Although a clear and consistent message is hard to extract, a few general trends seem to emerge. First, there is a tendency of results to be more similar for different methods with the same input mutation set, in particular for BRCA, where results for PyClone, SciClone, PhyloWGS and CSR are grouped together for each input set. Second, the really unexpected result is to observe that ITH results with the same input can be uncorrelated, and even significantly negatively correlated. Third, we observe two groups of methods that remain more similar across all three cancer types: EXPANDS and MATH score on the one hand, and PhyloWGS, PyClone, SciClone on the other hand. Those results can be related to the methods themselves. Indeed, PyClone, PhyloWGS and SciClone all define a probabilistic model explaining all observations of copy-number and read counts, based on a mixture model. They differ by the exact nature of the model (choice of distributions, exact definition of parameters), but they have similar structures. SciClone is different from PyClone and PhyloWGS in two ways: it only relies on mutations that are in regions without copy number alterations; and it does not correct for tumor purity. It is therefore not surprising that PyClone and PhyloWGS yield similar results, and that SciClone is a bit more different. CSR performs a consensus of all obtained clusterings; since 3 methods out of 4 have similar results, CSR might be biased towards those 3 methods. Expands makes similar assumptions as PyClone and PhyloWGS. However, the estimation process is very different: Expands estimates a distribution of read number for each position, and then clusters those distributions, while PyClone, SciClone and PhyloWGS attempt to find a common distribution for a group of mutations. The MATH score has an entirely different rationale as it simply ignores CNVs.Similar trends are observed when comparing the methods based on other pairwise comparison metrics (see Supplementary Figure S2-S4).

Regarding potential confounding variables, previous studies have reported a correlation between MATH score and CNA abundance [53, 22, 54], or between purity and ITH, as ITH methods were initially designed to refine purity estimation [29], and we observe similar behaviors. Association with immune infiltration has also been considered [54], though it is worth noting that immune infiltration and tumor purity are not independent, as immune cells are not cancerous. Each group of ITH methods is highly correlated to distinct genomic metrics, mutation load and CNV abundance (perc non diploid) for the first group (MATH, Expands), and purity (and the opposite of immune cells infiltration (inv T cells)) for the latter (PyClone, SciClone, PhyloWGS CSR). This might be indicative of systematic biases in the different methods, rather than biological strong signal as previously reported. Indeed, the strength and direction of all correlations vary between the two groups of ITH methods, and is hence hardly reliable or interpretable in terms of clinically actionable information without more data.

Similar results are obtained on an independent dataset of 7 samples from 5 patients where both WES and single cell sequencing was performed. In this dataset, subclonal reconstruction was performed by the method B-SCITE [42] that uses both bulk sequencing and single cell sequencing as input, and provides the most accurate representation possible. To further illustrate the behavior of ITH methods, we have compared results obtained for each sample separately to the B-SCITE result. To evaluate the concordance of each reconstruction to the B-SCITE reconstruction, we compare the number of clones, and the score2A from [40] that evaluates the co-clustering of mutations. The other metrics considered in [40] focus on the distance between the true and reconstructed cancer cell fraction distributions (score 1C), but in this setting, the ground truth does not provide a true CCF distribution estimate, and on the phylogenetic relationships between clones (score 3), but only PhyloWGS provides a tree among the considered methods. For this evaluation, we have left the true estimate for PyClone that provides a lot (several dozens) of clones with a single mutation. The input to ITH methods we have used results from variant calling on the bulk WES data, whereas the input to B-SCITE is more restrictive, and focuses on mutations detected both in the WES and in the single cells; the score2A is computed on the common mutations. Results are presented in Table 2. As observed on the TCGA, different methods based on WES data exhibit very different estimates of the number of clones, and none is very close to the estimates of B-SCITE using WES and single-cell data. In terms of clone composition, PyClone is the closest to B-SCITE in terms of score2A correlation in four out or seven samples, although the score2A values remain very modest.

### ITH is a weak and non robust prognosis factor

To test the prognostic power of each ITH quantification method, we collected survival information for the 686 patients on which all ITH methods ran successfully, and assessed how each ITH method allows to predict survival. Since all ITH methods except MATH output several features related to ITH, we did not test each feature individually but instead estimated a combined score for each method with a survival SVM model (see Methods). More precisely, we extract 5 features from each ITH method: the number of subclonal populations, the proportion of SNVs that belong the the major clone, the minimal cellular prevalence of a subclone, the Shannon index of the clonal distribution, and the cellular prevalence of the largest clone in terms of number of SNVs that enable to distinguish several evolutionary patterns, like early (star-like evolution) or late (tree with a long trunk) clonal diversification (see Methods). We evaluate the performance of each score by 5-fold cross-validation, and prognostic power is assessed on the test fold by computing the concordance index between the SVM prediction and the true patient survival. For MATH, a single feature is computed, so this procedure simply evaluates the concordance index of the MATH score with survival. In addition, we consider a model where all features of all methods (i.e., a total of 6 x 5 + 1 = 26 features) are combined together.

Figure 5 shows the results for each cancer type, each method, and each set of mutations used.

**Figure 5:**
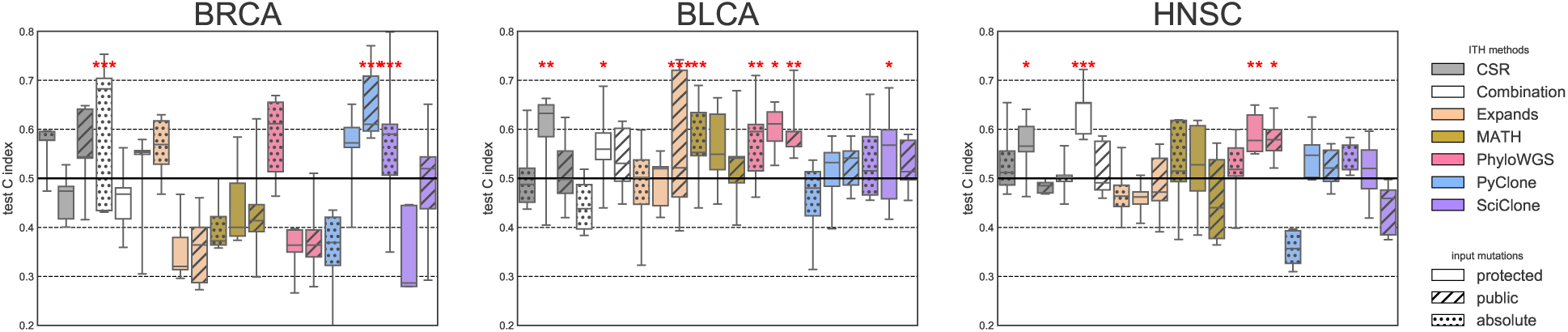
Prognostic power of ITH measured using different ITH method and input mutation set combination. 0.5 corresponds to a random prediction, and stars indicate statistical significance (p-value *<* 0.001: ***, *<* 0.01: **, *<* 0.05: *). Results are presented for 3 cancer types (BRCA, BLCA and HNSC from left to right).

Overall, we observe at least one method achieving significant survival prediction in each cancer type. The combined model is significantly prognostic with both protected and public sets in all three cancer types. Among the three cancer types, in the best case, however, the median concordance index on the test sets barely reaches 0.6 (except with the combination with absolute copy number in BRCA, but with an important variance), which remains modest for any clinical use. This suggests that there may be a weak prognostic signal captured by ITH measurement, but it can not be observed consistently with a single method and a single variant and copy number calling pipeline in the three cancer types, illustrating the frailty of obtained results. The combined model seems to be a robust alternative, as when it is significant, it has a concordance index in the range of the best performing single method; however the case of BRCA seems particular, as many methods perform worse than random.

Some authors [14, 55] have suggested a non-linear relation between survival and ITH, as very high ITH might be damaging for the tumor, while moderate ITH would be associated with aggressive tumors and prone to treatment resistance. To test this hypothesis in our framework we added squared features to the survival model, allowing second order polynomial relations between ITH and survival. However, this did not significantly impact the results (Supplementary S1). Indeed, after multiple test correction, only PyClone with the protected mutation set and ABSOLUTE copy number in BRCA prognostic power is increased by adding the squared features (*p* = 0.027, paired t-test), but both CI indexes remain below 0.5. We also assessed whether the relatively poor performance of the different methods was due to the difficulty to learn a prognostic score combining 5 features from limited amounts of training samples, by assessing the prognostic ability of a single feature: the number of clones. A significant improvement was obtained for 7and a significant decrease in performance in 3 of the 36 tested settings (4 methods, 2 mutation sets, 2 copy number methods, 3 cancer types). This suggests that the complexity of the model (polynomial of order 2 instead of linear) and the choice of ITH features have little influence on the results. This might be related to the fact that very little signal can be detected in the first place.

### ITH prognosis signal is redundant with other known factors

We have established that in some cases, ITH may exhibit weak prognostic power. It is then very important to assess whether it is complementary to already available prognostic features, like clinical characteristics. To answer this question, we consider relevant clinical features, as described in Methods.

Figure 6 presents a comparison between different prediction settings: clinical features without any clonality and clonality associated with clinical features. In all cases, clinical features alone have a significant prognostic power (median CI=0.79 for BRCA, 0.65 for BLCA, 0.65 for HNSC). More importantly, when we combine each ITH feature set with clinical features, we observe no significant improvement over clinical features alone. This suggests that the weak prognostic signal captured by ITH measures is in fact redundant with already available clinical factors.

**Figure 6:**
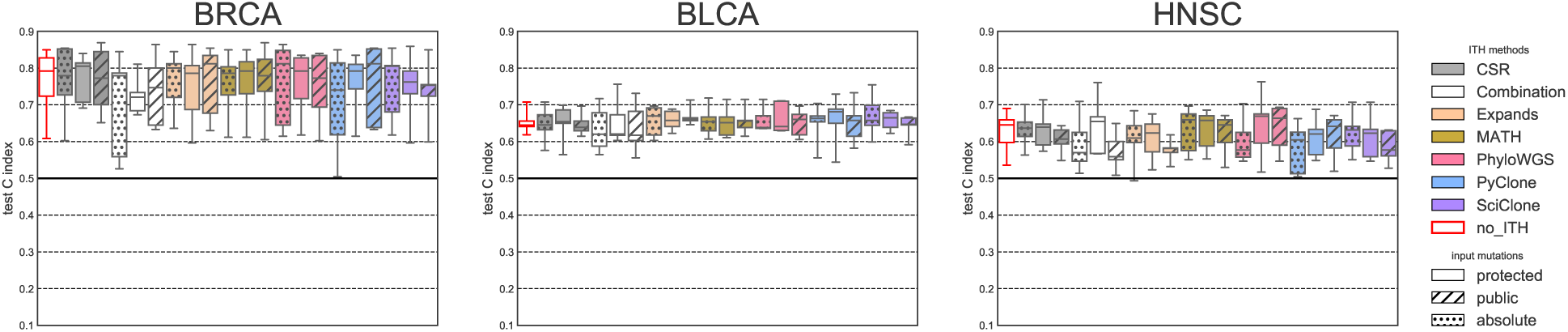
Prognostic power of ITH-derived features compared to other prognostic factors (686 patients in total). ITH-derived features are used in association with clinical features to predict overall survival. Left-most boxplots (with red contour lines) represent results using clinical variables alone, without any ITH, to serve as reference.

## Discussion

### Comparison to similar studies

Previous findings report divergent prognostic power for ITH in several pan cancer studies [14, 13, 22]. Andor et al [14] analyzed 1,165 patients across 12 cancer types from the TCGA, and found an overall prognostic power by considering all types together, and suggested that this effect might be nonlinear, with a trade-off between ITH and overall survival [55]. However, the association between the number of subclones and overall survival was significant with EXPANDS, but not with PyClone results, and no significant association was detected when considering each cancer type separately, except for gliomas. This might be due to the small number of cases of each type (between 33 and 166). Morris et al. [13] considered 3,383 patients of 9 cancer types from the TCGA and found significant association between the number of subclones found by PyClone in 5 types: HNSC, BRCA, KIRC, LGG, and PRAD. Noorbakhsh et al. [22] studied 4,722 patients from 11 types from the TCGA, and found significant prognostic power in 4 types using MATH score and distinct input mutation sets from different variant callers. They obtain significant prognostic association for all variant calling results in only one cancer type: UCEC, and already report some lack of robustness in the results. We have been further in testing up to 7 ITH methods with 2 alternative input mutation sets, in addition to the combination of all methods, and found no significant association, either for the same framework in all considered cancer types, nor for the same cancer type with all frameworks. We have also tested more powerful polynomial models to account for a potential nonlinear relationship, and results were inconclusive. This is an important distinction, because mutation calling can be made robust by additional experiments (targeted sequencing on WXS or WGS candidates), but our results highlight intrinsic limitations of ITH methods.

Considering results in details, there are discrepancies that should be discussed. For BRCA, conclusions are more discordant: Morris et al. [13] found significant results, Noorbakhsh et al. [22] did not, and in more specialized studies like METABRIC [53], significant association was found when considering only the upper and lower quartile of MATH score for ER+ tumors. For BLCA, contradictory conclusions were also drawn, as previous studies [14, 13] found no prognostic power and we have with some ITH methods. There are several explanations: each study considered a distinct subset of patients, with a distinct pipeline for calling mutations and measure ITH. This instability with respect to patient selection has been confirmed by our study. All of those studies, including ours, observed ITH prognostic relevance in HNSC. Good prognostic power for HNSC and BLCA might be an indication that the importance of ITH for cancer aetiology differs across cancer types.

### Can we truly measure ITH?

Beyond the question of the prognostic power of ITH, our results challenge the very fact that ITH can be measured accurately with one WES sample per patient. Up to 30 methods have been developed to tackle ITH detection and quantification from NGS data in tumor samples, and new ones are still being developed [56]. This analysis has focused on relatively early but among the most widely used ITH methods in order to provide valuable insight on the degree of reliability of provided results. Indeed results presented here show that there is a very weak correlation (and sometimes even a significant negative correlation) between results obtained with different methods on the same patients. Another source of inconsistency is that ITH methods rely on results from previous analysis steps, in particular variant calling. Indeed, all ITH methods rely on the distribution of SNV frequencies, in association or not with structural variants (also called by a variety of dedicated methods). This has already been discussed by Noorbakhsh et al. [22] for MATH score computation. We show here that this issue is not limited to the MATH score. Some authors have suggested that being very restrictive in variant calling, even resorting to targeted deep sequencing to experimentally validate SNVs [6], would exhibit less noisy results. Here we have not observed any evidence that ITH methods estimated more robust results with a restricted input mutation set (i.e. the public mutation set in this study). Overall, lack of agreement between the different ITH measures is a real concern, indicating again that ITH is probably not very accurate. A similar conclusion was recently and independently reached by [57].

Beyond the methods used for ITH inference, the data might also be questioned. Being able to measure ITH to one sample WES with moderate sequencing depth is tempting for future clinical application where the cost and the inconvenience of multiple samples for patients should be limited [3], but it may be unrealistic, as the true heterogeneity of a tumor can be missed by a single biopsy. However, more complex experimental settings have allowed more convincing findings in the field of tumor evolution [58, 59], and it may be necessary to further evaluate lack of accuracy due to undersampling from the whole tumor or to use of WES instead of WGS, and the impact of sequencing depth. A recent and broad analysis of ITH with one WGS sample per patient [16] partially answers as the authors could detect ITH in almost every patient, and conduct interesting further analyses as they had confidence in the robustness of ITH estimates. Most published methods are able to account for multiple samples from the same patient, either sampled at different times or from different regions of the tumor. However, for extension to WGS analysis, our work highlights limitations with respect to the computation time for high numbers of mutation as input.

### Association with survival, link with other variables

It is tempting to formulate the hypothesis that higher association with patient survival is a sign of higher accuracy. We have already mentioned some technical issues associated with the setting of one sample WES per patient, as even without measure issues, ITH might just be under-represented in the sample compared to the whole tumor [60]. Another limitation is that this does not represent a dynamic measure. For instance a tumor can be clonal because it is not very aggressive, or on the contrary this might be the result of a selective sweep after a phase of new clonal expansion. Moreover, several authors discuss the consequences and the interplay of the presence of distinct subclonal populations, in terms of cooperation [61, 62], competition [63, 64], or even neutral evolution [65, 66]. Hence, the same level of ITH might uncover very diverse situations, and may not be a prognostic factor by itself.

Moreover, the dataset used in this survival analysis has some particularities: the TCGA has selected patients with criteria allowing high sequencing quality, and ITH analysis itself has further eliminated tumors with no or very high CNA abundance, which may also bias results. Finally, absence of prognosis power in one dataset does not constitute a formal proof that ITH is not associated with survival.

Besides, ITH is likely to be influenced and to interplay with other external factors including tumor micro-environment, immune response, nutrient availability. Recent work has tried to set a full framework for analysis including many factors [67]. However, in the case of the TCGA, not all those variables are measurable, but some might be included in further work. In this line of thought, earlier results exhibited correlation of ITH with other factors like CNA abundance, sample purity, immune infiltration [53, 54, 68, 13]. Our results show that the strength (and even direction in the case of CNA abundance and mutation load) of correlation between those factors and ITH varies between the different tested ITH measures. This again calls for further and more detailed analysis, as results show ambiguity and lack of robustness.

### Can we build a gold standard dataset for benchmark?

The main difficulty of ITH estimation is to assess the accuracy of the results. In this work, we have considered two possibilities. The first one on data from the TCGA is to work without any ground truth proxy and measure other features of accuracy: robustness, agreement of results obtained by different methods and association with other clinical variables. The obtained results suggest that the considered ITH methods are relatively robust to changes in the copy number input, but very sensitive to the input mutations. The last two options are more difficult to work with, as one method could be in disagreement with all the others but still provide the most accurate result, and absence or presence of association between ITH and other clinical or genomic variables can be either due to a real biological signal or be an artifact (or bias) of the method. Though the goal of this study is not to provide a formal evaluation of the considered method, the results on the TCGA provide information on systematic trends of each method, and the level of confidence to expect when applying ITH methods.

A second possibility is to try and obtain a proxy for the ground truth. This can be done using single cell sequencing in addition to the bulk sequencing. Though suffering from other issues, single cell sequencing provides true associations or exclusions of mutations, and hence constraints the subclonal reconstruction [42]. However, a large number of cells is necessary. In the 7 samples considered in this study, only a subset of the mutations identified in the bulk sequencing were also identified in single cells, limiting the representativity and the relevance of the extracted accuracy measures. A second possibility is to rely on several samples from the same tumor to obtain a better ground truth to compare to the result obtained with one sample. However, each sample is a priori heterogeneous itself, requiring a first multi-sample deconvolution. This first step can be challenging, as it is thought that multi-sample reconstruction is subject to a larger statistical bias compared to single sample reconstruction [69], and the accuracy of this first step will be critical in the final results. A final possibility is to rely on simulated data, which have the major drawback to not be necessarily representative of the true biological data, as recently highlighted for ITH in [69], that point to an aspect of the input data so far overlooked by the community.

## Acknowledgments

The authors declare no potential conflicts of interest. We thank Peter Van Loo and Kerstin Haase for sharing ASCAT results on the TCGA, and Christoffer Flensburg for his help with the liftover of ASCAT results. We thank the authors of ”Integrative inference of subclonal tumor evolution from single-cell and bulk sequencing data” for help in adapting their results and reprocess the data, and Elodie Girard for her help with NGS data processing. We also thank Alice Schoenauer Sebag for helpful discussions. The results shown here are based upon data generated by the TCGA Research Network: http://cancergenome.nih.gov/, under authorization for project 10569 (dbGaP).

## Supplementary information

**Figure S1:**
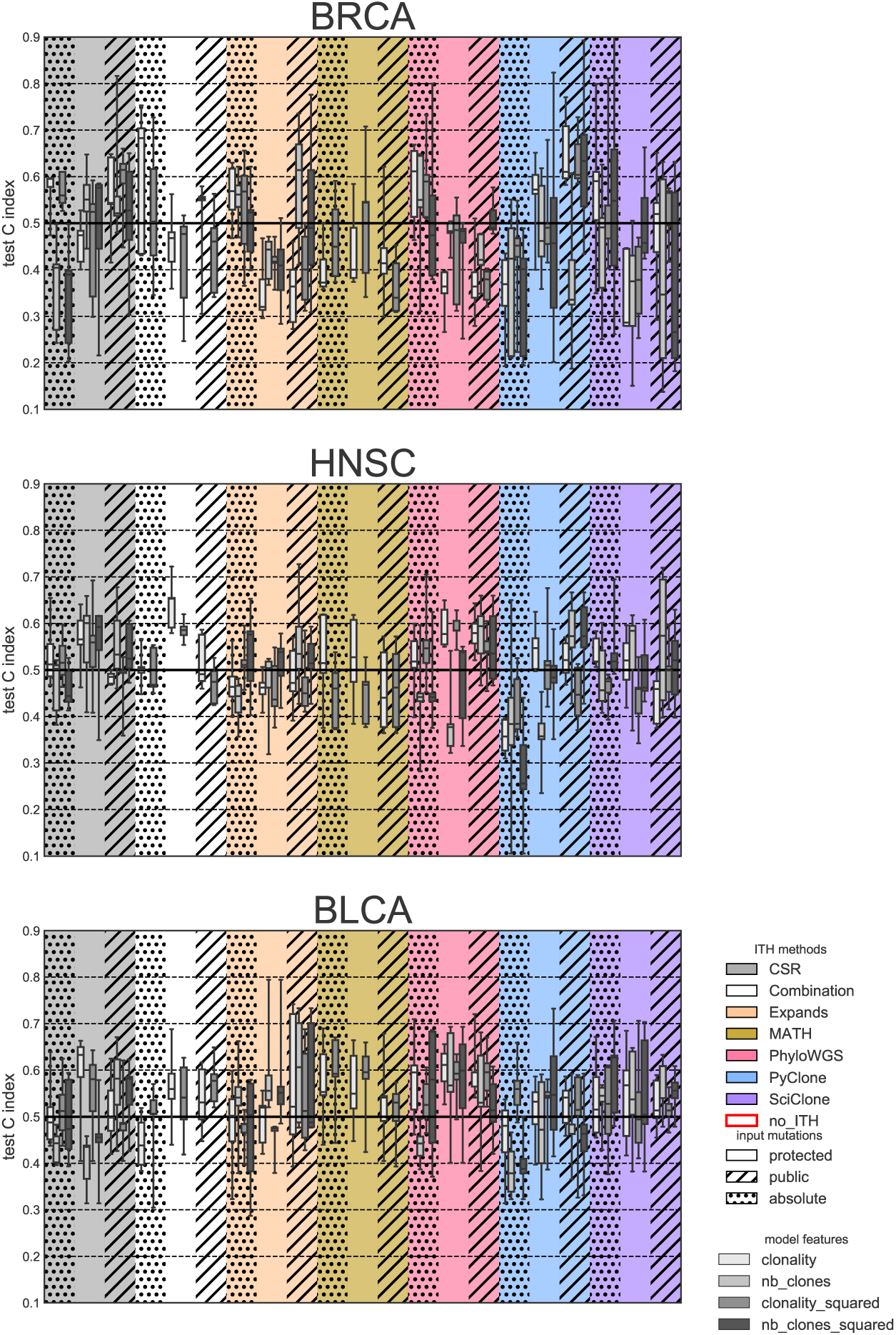
Prognostic power of diverse combination of ITH-derived features, on the three cancer types (respectively BRCA, HNSC and BLCA from top to bottom). In each plot, the background color indicates the ITH method used. Each method is tested on protected or public mutations (hashed). For each method, we assess the ability to predict survival with a survival SVM using 4 sets of features: (i) the number of clones alone, (ii) the five custom features which include the number of clones, and (iii) and (iv) the concatenations of features in (i) and (ii) with their squares, to account for possible nonlinear quadratic effects. We observe no clear trend of one of the two sets performs systematically better than the other, and the squared features have not significantly improved results either.

**Figure S2:**
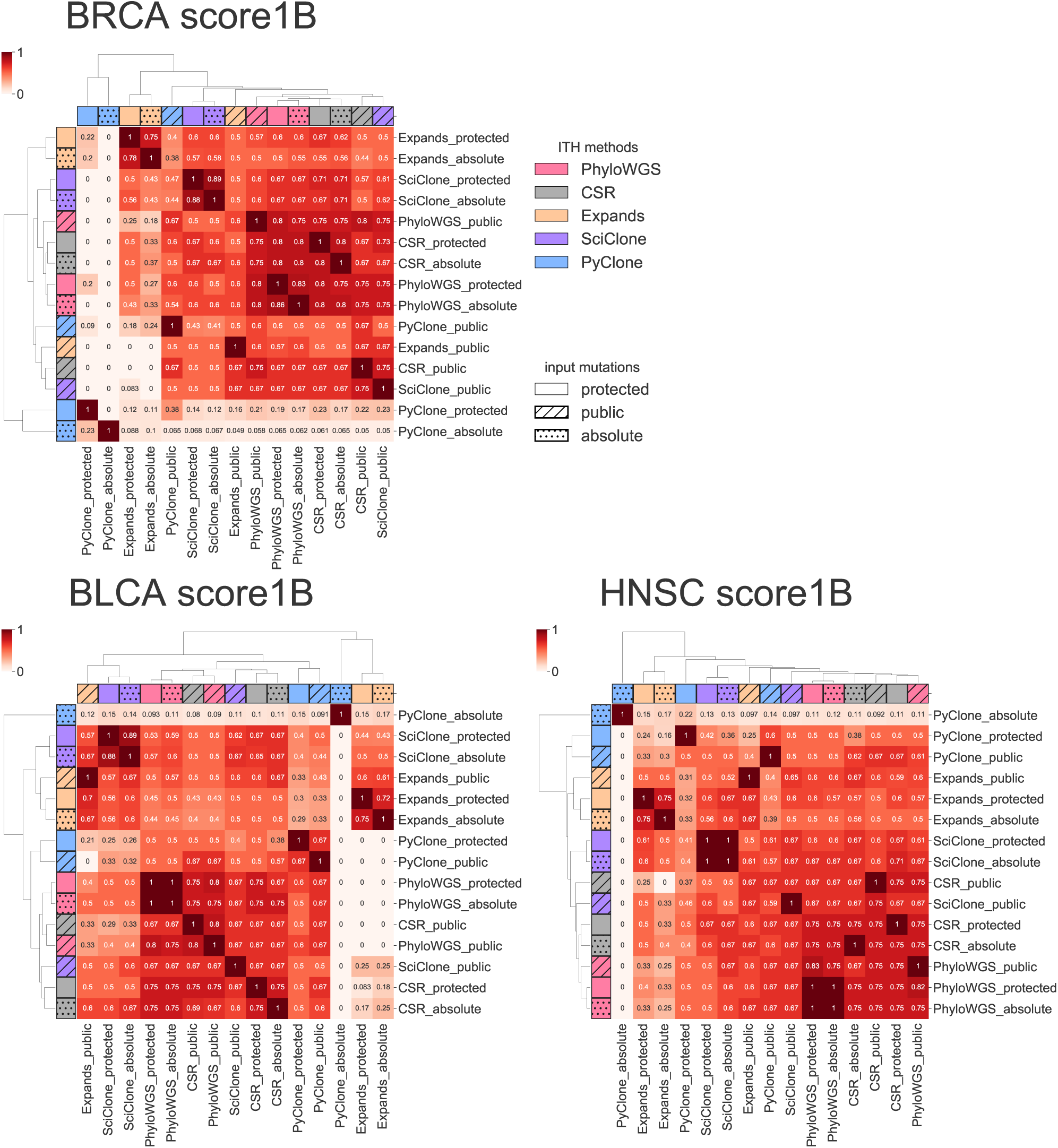
Pairwise computation of score1B for the different ITH methods and inputs. Score1B is a metric designed in [40] penalizes differences between the number of clones inferred in each case in a symmetric way (only the difference matters, either more or fewer clones are detected), following the formula 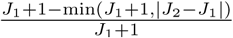, with *J*_1_ and *J*_2_ the numbers of clones found by each method. The score was computed for all patients, and this heatmap represents the median score. We observe a particular feature of PyClone, which tends to find a lot (sometimes several dozens) of clones with only one mutation. They were discarded when comparing the number of clones, but not for the computation of metric 1B to ensure consistency with the other metrics.

**Figure S3:**
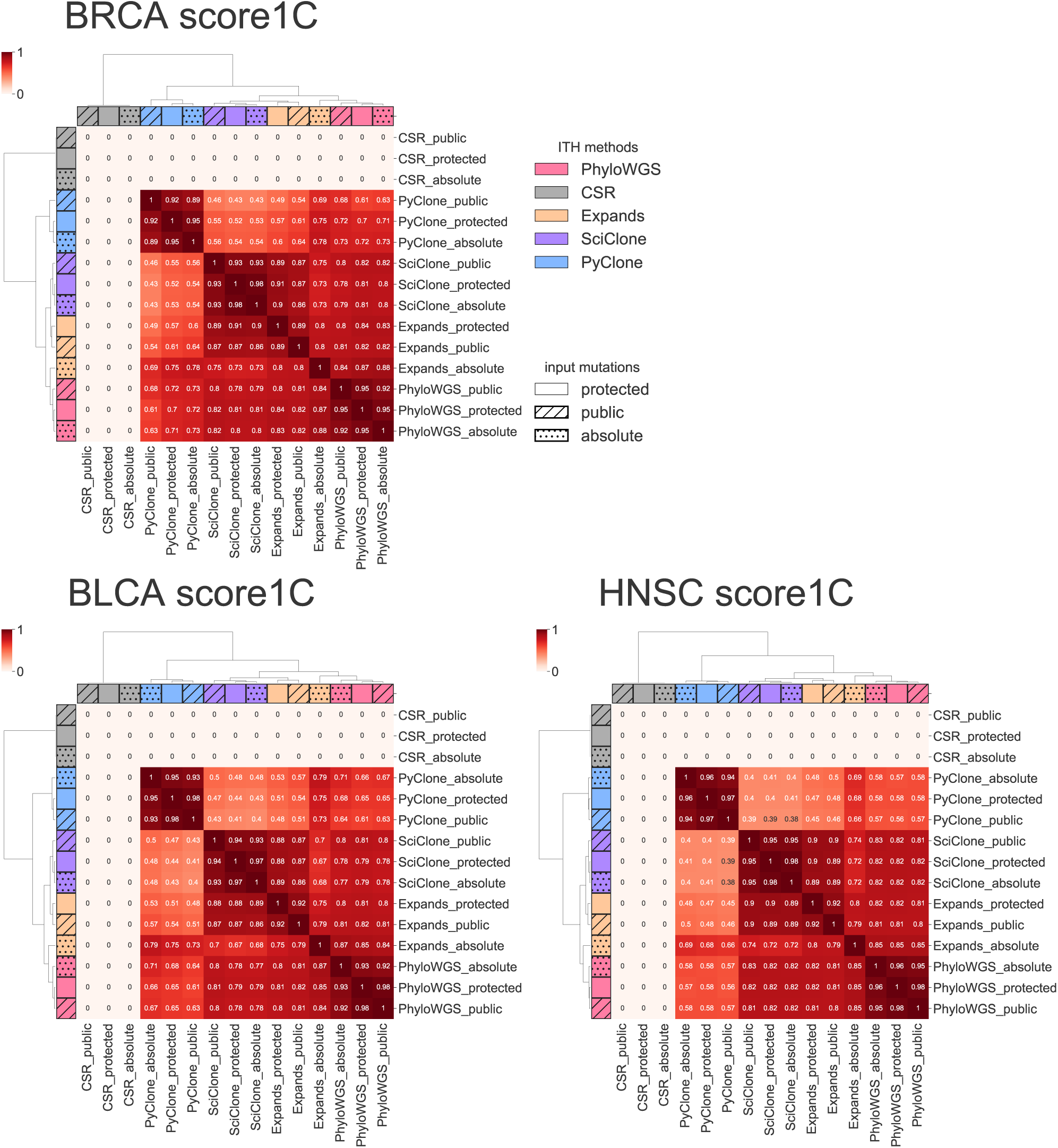
Pairwise computation of score1C for the different ITH methods and inputs. Score1C is a metric designed in [40] that represents the Wasserstein distance between the cancer cell fraction (CCF) distribution resulting from each clone’s mean CCF and number of mutations. Due to the number of single-mutation clones of PyClone, the resulting distribution is quite different from the other cases. As CSR only takes as input the mutation attribution to clones by other methods, without taking into account their CCF, we did not compute score1C for that method. The score was computed for all patients, and this heatmap represents the median score.

**Figure S4:**
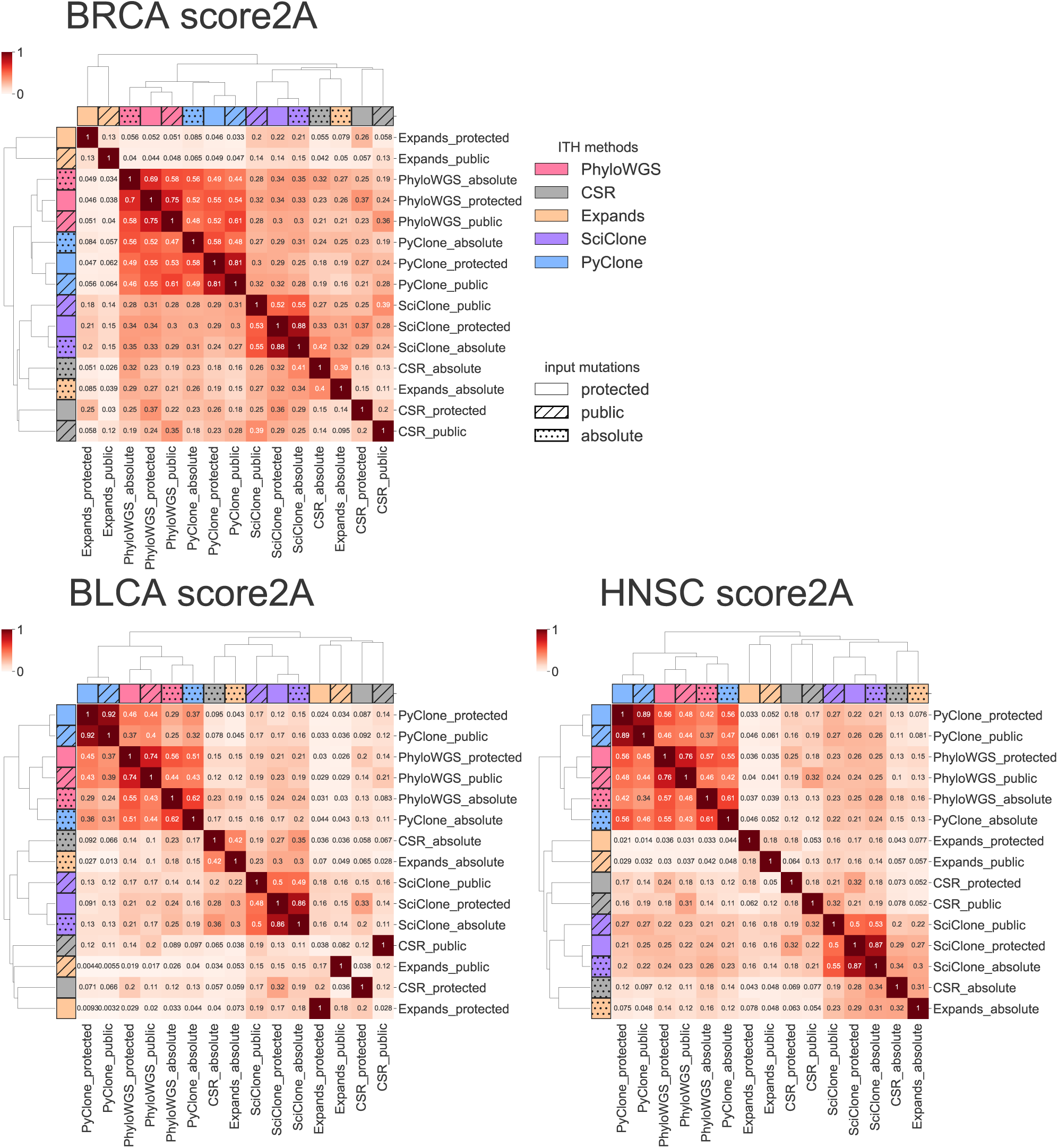
Pairwise computation of score2A for the different ITH methods and inputs. Score2A is a metric designed in [40] that assesses the similarity of the mutation clustering resulting from subclonal reconstruction (see Methods for details). We recover the previously observed pattern that PyClone and PhyloWGS are the closest methods. The score was computed for all patients, and this heatmap represents the median score.

**Table S1:** Summary statistics of the number of protected and public mutations per sample for BRCA, BLCA and HNSC samples. The protected set corresponds to raw variant calling outputs. The public set corresponds to publicly available SNV calls.

|  | BRCA | BLCA | HNSC |
| --- | --- | --- | --- |
| number of samples | 962 | 351 | 445 |
| **Protected mutations** |  |  |  |
| average | 530 | 735 | 456 |
| std | 1,189 | 890 | 544 |
| min | 79 | 61 | 31 |
| median | 342 | 525 | 340 |
| max | 21,821 | 12,774 | 7,941 |
| **Public mutations** |  |  |  |
| average | 121 | 352 | 202 |
| std | 375 | 426 | 271 |
| min | 1 | 2 | 1 |
| median | 63 | 241 | 140 |
| max | 7,919 | 5,478 | 3,935 |

**Table S2:**
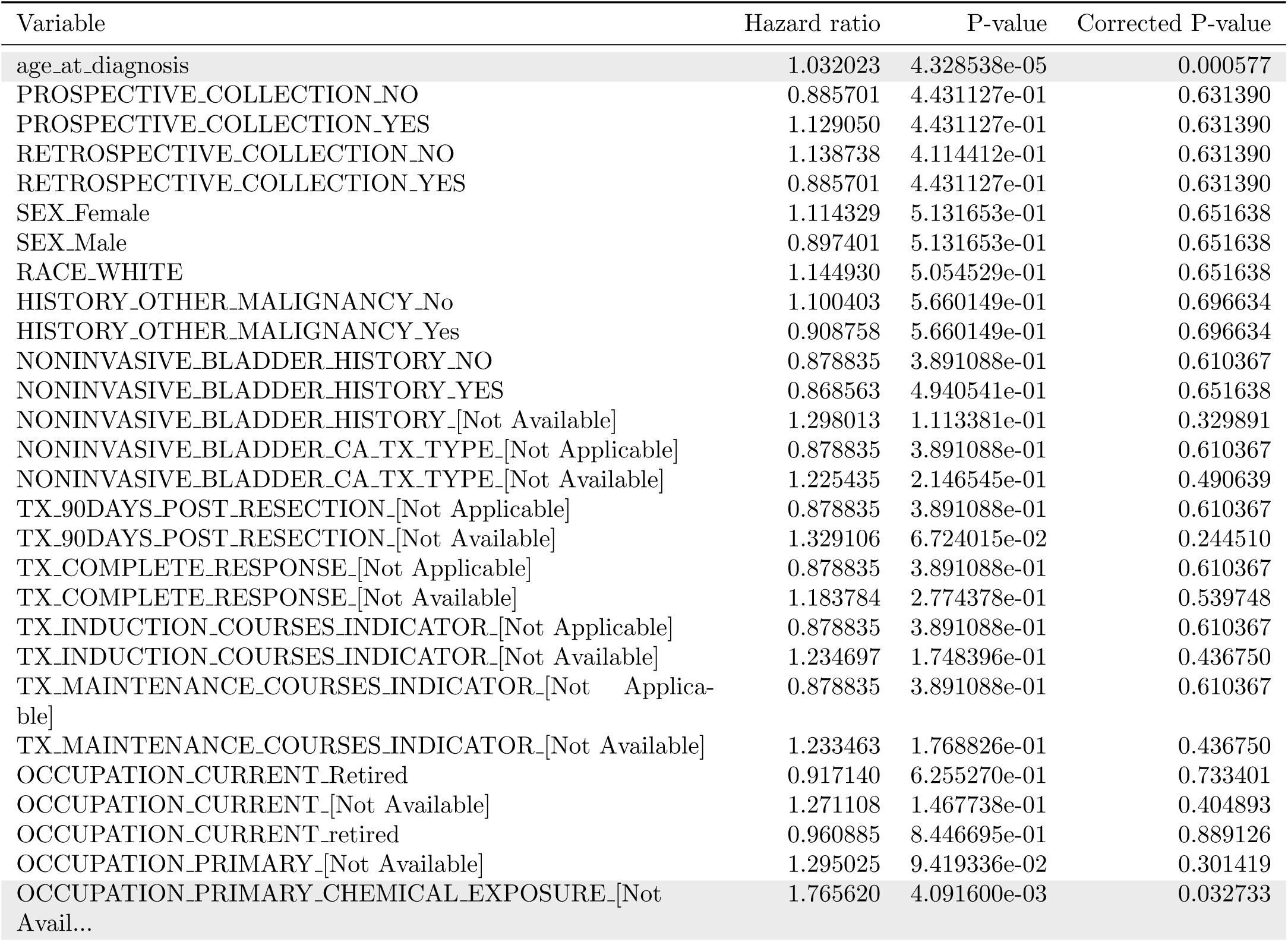
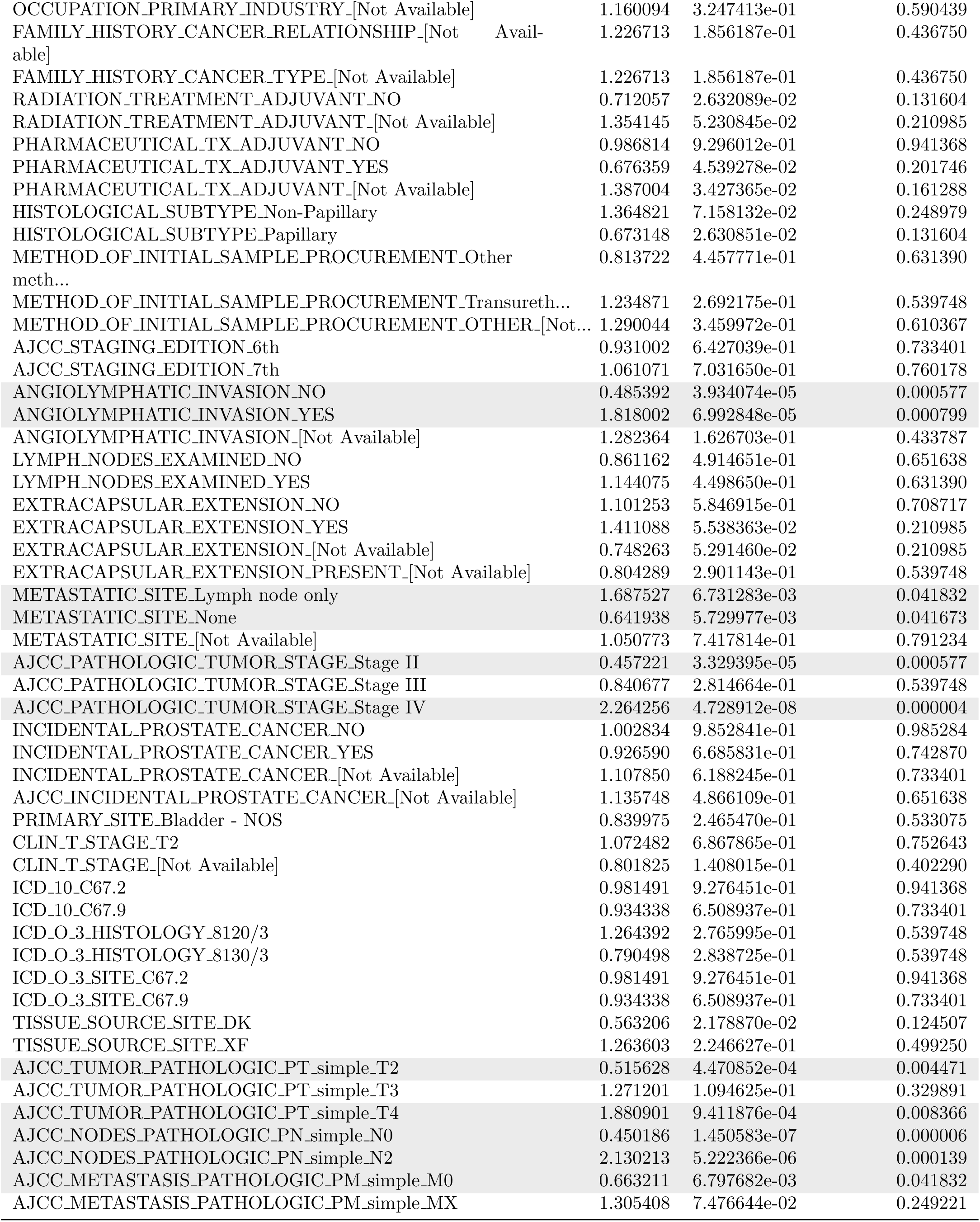
Clinical variables significance for single-variable cox model for BLCA (409 patients). Variable significantly associated with survival are shaded.

**Table S3:** Clinical variables significance for single-variable cox model for BRCA (1080 patients). Variable significantly associated with survival are shaded.

| Variable | Hazard ratio | P-value | Corrected P-value |
| --- | --- | --- | --- |
| age_at_diagnosis | 1.032118 | 4.566898e-07 | 1.077788e-05 |
| PROSPECTIVE_COLLECTION_NO | 0.602422 | 4.530180e-02 | 1.243166e-01 |
| PROSPECTIVE_COLLECTION_YES | 1.686136 | 3.888498e-02 | 1.176520e-01 |
| RETROSPECTIVE_COLLECTION_NO | 1.686136 | 3.888498e-02 | 1.176520e-01 |
| RETROSPECTIVE_COLLECTION_YES | 0.602422 | 4.530180e-02 | 1.243166e-01 |
| MENOPAUSE_STATUS_Post (prior bilateral ovariect... | 1.324224 | 9.572180e-02 | 2.026486e-01 |
| MENOPAUSE_STATUS_Pre (<6 months since LMP AND n... | 0.431889 | 8.809889e-04 | 5.887468e-03 |
| MENOPAUSE_STATUS_[Not Available] | 2.943808 | 9.817809e-07 | 1.930836e-05 |
| RACE_BLACK OR AFRICAN AMERICAN | 1.215466 | 3.390936e-01 | 5.275821e-01 |
| RACE_WHITE | 0.825045 | 2.987240e-01 | 4.828689e-01 |
| RACE_[Not Available] | 1.274439 | 5.361448e-01 | 6.741716e-01 |
| ETHNICITY_NOT HISPANIC OR LATINO | 2.207632 | 1.176302e-02 | 5.552148e-02 |
| ETHNICITY_[Not Available] | 0.603636 | 1.259999e-01 | 2.447704e-01 |
| HISTORY_OTHER_MALIGNANCY_No | 0.669493 | 2.469654e-01 | 4.047489e-01 |
| HISTORY_OTHER_MALIGNANCY_Yes | 1.501028 | 2.411757e-01 | 4.016651e-01 |
| RADIATION_TREATMENT_ADJUVANT_NO | 0.825072 | 6.454644e-01 | 7.693415e-01 |
| RADIATION_TREATMENT_ADJUVANT_YES | 1.184551 | 6.240900e-01 | 7.525332e-01 |
| RADIATION_TREATMENT_ADJUVANT_[Not Available] | 0.991267 | 9.744419e-01 | 9.936324e-01 |
| PHARMACEUTICAL_TX_ADJUVANT_YES | 0.893189 | 7.194462e-01 | 8.162947e-01 |
| PHARMACEUTICAL_TX_ADJUVANT_[Not Available] | 1.001332 | 9.961223e-01 | 9.961223e-01 |
| METHOD_OF_INITIAL_SAMPLE_PROCUREMENT_Core<br>needl... | 0.545734 | 3.371119e-04 | 2.841372e-03 |
| METHOD_OF_INITIAL_SAMPLE_PROCUREMENT_Fine<br>needl... | 1.669895 | 1.358907e-02 | 5.938927e-02 |
| METHOD_OF_INITIAL_SAMPLE_PROCUREMENT_Other<br>meth... | 1.009070 | 9.765300e-01 | 9.936324e-01 |
| METHOD_OF_INITIAL_SAMPLE_PROCUREMENT_Tumor<br>rese... | 0.740284 | 3.397987e-01 | 5.275821e-01 |
| METHOD_OF_INITIAL_SAMPLE_PROCUREMENT_[Not<br>Avail... | 1.944929 | 3.630026e-02 | 1.157684e-01 |
| METHOD_OF_INITIAL_SAMPLE_PROCUREMENT_OTHER_[Not... | 0.994946 | 9.868232e-01 | 9.952576e-01 |
| SURGICAL_PROCEDURE_FIRST_Lumpectomy | 0.690083 | 9.704508e-02 | 2.026486e-01 |
| SURGICAL_PROCEDURE_FIRST_Modified Radical Maste... | 1.476196 | 2.170723e-02 | 8.004541e-02 |
| SURGICAL_PROCEDURE_FIRST_Other | 0.882243 | 5.027363e-01 | 6.448140e-01 |
| SURGICAL_PROCEDURE_FIRST_Simple Mastectomy | 0.657380 | 9.788958e-02 | 2.026486e-01 |
| SURGICAL_PROCEDURE_FIRST_[Not Available] | 2.480055 | 2.729308e-03 | 1.463901e-02 |
| FIRST_SURGICAL_PROCEDURE_OTHER_Surgical Resection | 1.534372 | 3.107861e-01 | 4.955778e-01 |
| FIRST_SURGICAL_PROCEDURE_OTHER_[Not Available] | 1.084261 | 6.601793e-01 | 7.790115e-01 |
| PATH_MARGIN_Negative | 0.328745 | 2.487080e-11 | 1.467377e-09 |
| PATH_MARGIN_Positive | 1.456750 | 1.265338e-01 | 2.447704e-01 |
| PATH_MARGIN_[Not Available] | 4.295169 | 1.280242e-14 | 1.510685e-12 |
| SURGERY_FOR_POSITIVE_MARGINS_[Not Available] | 0.881873 | 7.014933e-01 | 8.141488e-01 |
| MARGIN_STATUS_REEXCISION_Negative | 1.189978 | 5.498957e-01 | 6.830283e-01 |
| MARGIN_STATUS_REEXCISION_[Not Available] | 0.788031 | 3.970790e-01 | 5.714063e-01 |
| STAGING_SYSTEM_Axillary lymph node dissection a... | 1.147628 | 4.478192e-01 | 6.047185e-01 |
| STAGING_SYSTEM_Sentinel lymph node biopsy plus ... | 0.471890 | 9.803200e-04 | 6.088303e-03 |
| STAGING_SYSTEM_Sentinel node biopsy alone | 0.517762 | 9.212199e-03 | 4.529331e-02 |
| STAGING_SYSTEM_[Not Available] | 2.306054 | 2.789049e-06 | 3.502342e-05 |
| MICROMET_DETECTION_BY_IHC_NO | 0.983191 | 9.205373e-01 | 9.860025e-01 |
| MICROMET_DETECTION_BY_IHC_YES | 0.330809 | 1.172534e-05 | 1.257809e-04 |
| MICROMET_DETECTION_BY_IHC_[Not Available] | 2.187550 | 2.968086e-06 | 3.502342e-05 |
| LYMPH_NODES_EXAMINED_YES | 1.127012 | 4.842736e-01 | 6.279592e-01 |
| LYMPH.NODES_EXAMINED_[Not Available] | 0.789955 | 1.767459e-01 | 3.160002e-01 |
| AJCC_STAGING_EDITION_5th | 2.522972 | 2.425021e-06 | 3.502342e-05 |
| AJCC_STAGING_EDITION_6th | 0.391254 | 3.279076e-07 | 9.673274e-06 |
| AJCC_STAGING_EDITION_7th | 1.553473 | 5.971021e-02 | 1.455199e-01 |
| AJCC_STAGING_EDITION_[Not Available] | 0.905993 | 7.048542e-01 | 8.141488e-01 |
| AJCC_PATHOLOGIC_TUMOR_STAGE_Stage I | 0.623674 | 1.050290e-01 | 2.136796e-01 |
| AJCC_PATHOLOGIC_TUMOR_STAGE_Stage IA | 0.246046 | 1.650590e-02 | 6.956059e-02 |
| AJCC_PATHOLOGIC_TUMOR_STAGE_Stage IIA | 0.697084 | 6.472348e-02 | 1.510602e-01 |
| AJCC_PATHOLOGIC_TUMOR_STAGE_Stage IIB | 0.828285 | 3.501038e-01 | 5.365227e-01 |
| AJCC_PATHOLOGIC_TUMOR_STAGE_Stage IIIA | 1.293391 | 2.416799e-01 | 4.016651e-01 |
| AJCC_PATHOLOGIC_TUMOR_STAGE_Stage IIIC | 1.898479 | 6.528872e-02 | 1.510602e-01 |
| ER_STATUS_BY_IHC_Negative | 1.272320 | 1.906432e-01 | 3.308221e-01 |
| ER_STATUS_BY_IHC_Positive | 0.662619 | 1.727163e-02 | 7.027765e-02 |
| ER_STATUS_IHC_PERCENT_POSITIVE_90-99% | 0.370064 | 2.554685e-03 | 1.435490e-02 |
| ER_STATUS_IHC_PERCENT_POSITIVE_≤10% | 0.771901 | 4.520339e-01 | 6.047185e-01 |
| ER_STATUS_IHC_PERCENT_POSITIVE_[Not Available] | 2.276830 | 1.936838e-05 | 1.758053e-04 |
| ER_POSITIVITY_SCALE_USED_3 Point Scale | 0.369684 | 5.031369e-02 | 1.349322e-01 |
| ER_POSITIVITY_SCALE_USED_[Not Available] | 2.435285 | 3.311614e-02 | 1.116487e-01 |
| ER_POSITIVITY_SCALE_OTHER_[Not Available] | 1.595260 | 1.827359e-01 | 3.218334e-01 |
| BRACHYTHERAPY_TOTAL_DOSE_POINT_A_[Not Available] | 0.994099 | 9.767911e-01 | 9.936324e-01 |
| PR_STATUS_BY_IHC_Negative | 1.257745 | 1.746637e-01 | 3.160002e-01 |
| PR_STATUS_BY_IHC_Positive | 0.717066 | 4.336782e-02 | 1.243166e-01 |
| PR_STATUS_IHC_PERCENT_POSITIVE_90-99% | 0.082244 | 1.281943e-02 | 5.818049e-02 |
| PR_STATUS_IHC_PERCENT_POSITIVE_≤10% | 0.987074 | 9.609962e-01 | 9.936324e-01 |
| PR_STATUS_IHC_PERCENT_POSITIVE_[Not Available] | 2.440652 | 1.832734e-05 | 1.758053e-04 |
| PR_POSITIVITY_SCALE_USED_3 Point Scale | 0.401865 | 7.305251e-02 | 1.626452e-01 |
| PR_POSITIVITY_SCALE_USED_[Not Available] | 2.235079 | 5.428402e-02 | 1.411211e-01 |
| PR_POSITIVITY_IHC_INTENSITY_SCORE_3+ | 0.127355 | 4.012324e-02 | 1.183636e-01 |
| PR_POSITIVITY_IHC_INTENSITY_SCORE_[Not Available] | 3.001282 | 2.553857e-03 | 1.435490e-02 |
| PR_POSITIVITY_SCALE_OTHER_[Not Available] | 1.327000 | 3.966578e-01 | 5.714063e-01 |
| PR_POSITIVITY_DEFINE.METHOD_[Not Available] | 1.008404 | 9.676018e-01 | 9.936324e-01 |
| IHC.HER2_Equivocal | 0.601537 | 5.547120e-02 | 1.411211e-01 |
| IHC.HER2_Negative | 0.791443 | 1.711526e-01 | 3.155626e-01 |
| IHC.HER2_Positive | 1.504160 | 7.472326e-02 | 1.632842e-01 |
| IHC.HER2_[Not Available] | 1.460324 | 3.585853e-02 | 1.157684e-01 |
| HER2_IHC_PERCENT_POSITIVE_≤10% | 0.408510 | 2.110215e-02 | 8.004541e-02 |
| HER2_IHC_PERCENT_POSITIVE_[Not Available] | 1.864321 | 2.281919e-02 | 8.159588e-02 |
| HER2_POSITIVITY_METHOD.TEXT_[Not Available] | 2.593860 | 6.042777e-02 | 1.455199e-01 |
| HER2_FISH_STATUS_Negative | 0.739080 | 1.150908e-01 | 2.301816e-01 |
| HER2_FISH_STATUS_Positive | 0.880007 | 7.106553e-01 | 8.141488e-01 |
| HER2_FISH_STATUS_[Not Available] | 1.385428 | 6.801151e-02 | 1.543338e-01 |
| HER2_COPY_NUMBER_[Not Available] | 1.645350 | 1.487810e-01 | 2.831638e-01 |
| CENT17_COPY_NUMBER_[Not Available] | 1.548415 | 2.047798e-01 | 3.502032e-01 |
| PRIMARY_SITE_Left | 1.945749 | 7.591896e-04 | 5.599023e-03 |
| PRIMARY_SITE_Left Upper Inner Quadrant | 0.734898 | 4.268913e-01 | 6.047185e-01 |
| PRIMARY_SITE_Left Upper Outer Quadrant | 0.826630 | 4.397503e-01 | 6.047185e-01 |
| PRIMARY_SITE_Right | 0.892605 | 6.249852e-01 | 7.525332e-01 |
| PRIMARY_SITE_Right Upper Outer Quadrant | 1.138461 | 6.005181e-01 | 7.381368e-01 |
| HISTOLOGICAL_DIAGNOSIS_Infiltrating Ductal Carc... | 0.969817 | 8.637923e-01 | 9.351146e-01 |
| HISTOLOGICAL_DIAGNOSIS_Infiltrating Lobular Car... | 0.823077 | 3.912033e-01 | 5.714063e-01 |
| ICD.O.3_HISTOLOGY_8500/3 | 0.949261 | 7.692879e-01 | 8.563771e-01 |
| ICD.O.3_HISTOLOGY_8520/3 | 0.871304 | 5.370519e-01 | 6.741716e-01 |
| METASTATIC_TUMOR_INDICATOR_NO | 0.641384 | 2.830420e-02 | 9.823221e-02 |
| METASTATIC_TUMOR_INDICATOR_[Not Available] | 1.149966 | 4.502190e-01 | 6.047185e-01 |
| TISSUE_SOURCE_SITE_A2 | 0.816065 | 4.561012e-01 | 6.047185e-01 |
| TISSUE_SOURCE_SITE_A8 | 1.386201 | 4.396658e-01 | 6.047185e-01 |
| TISSUE_SOURCE_SITE_AR | 0.426279 | 7.036257e-03 | 3.609906e-02 |
| TISSUE_SOURCE_SITE_B6 | 1.053316 | 8.202897e-01 | 9.046185e-01 |
| TISSUE_SOURCE_SITE_BH | 2.772661 | 1.247824e-08 | 4.908109e-07 |
| TISSUE_SOURCE_SITE_D8 | 0.947428 | 9.275108e-01 | 9.860025e-01 |
| TISSUE_SOURCE_SITE_E2 | 0.555754 | 1.612908e-01 | 3.021003e-01 |
| TISSUE_SOURCE_SITE_E9 | 1.350339 | 4.798622e-01 | 6.279592e-01 |
| AJCC_TUMOR_PATHOLOGIC_PT_simple_T1 | 0.686002 | 5.620924e-02 | 1.411211e-01 |
| AJCC_TUMOR_PATHOLOGIC_PT_simple_T2 | 0.951778 | 7.620026e-01 | 8.563458e-01 |
| AJCC_TUMOR_PATHOLOGIC_PT_simple_T3 | 1.220149 | 3.652398e-01 | 5.525423e-01 |
| AJCC_NODES_PATHOLOGIC_PN_simple_N0 | 0.428539 | 2.290188e-06 | 3.502342e-05 |
| AJCC_NODES_PATHOLOGIC_PN_simple_N1 | 1.159871 | 3.741624e-01 | 5.588754e-01 |
| AJCC_NODES_PATHOLOGIC_PN_simple_N2 | 1.714021 | 1.986213e-02 | 7.812439e-02 |
| AJCC_NODES_PATHOLOGIC_PN_simple_N3 | 2.564601 | 6.195610e-04 | 4.873880e-03 |
| AJCC_METASTASIS_PATHOLOGIC_PM_simple_M0 | 0.509375 | 8.980883e-04 | 5.887468e-03 |
| AJCC_METASTASIS_PATHOLOGIC_PM_simple_MX | 1.062431 | 8.307274e-01 | 9.076466e-01 |

**Table S4:**
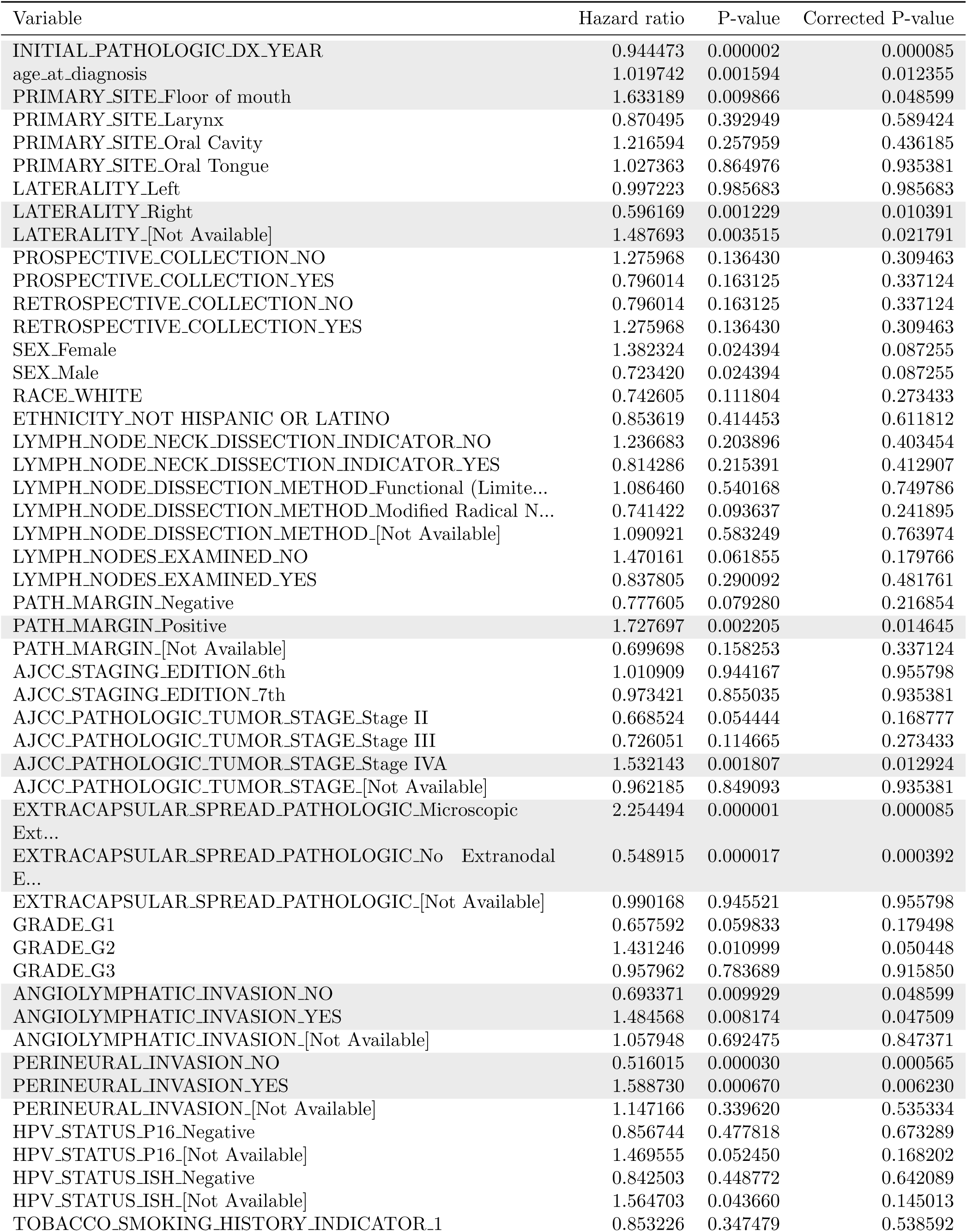
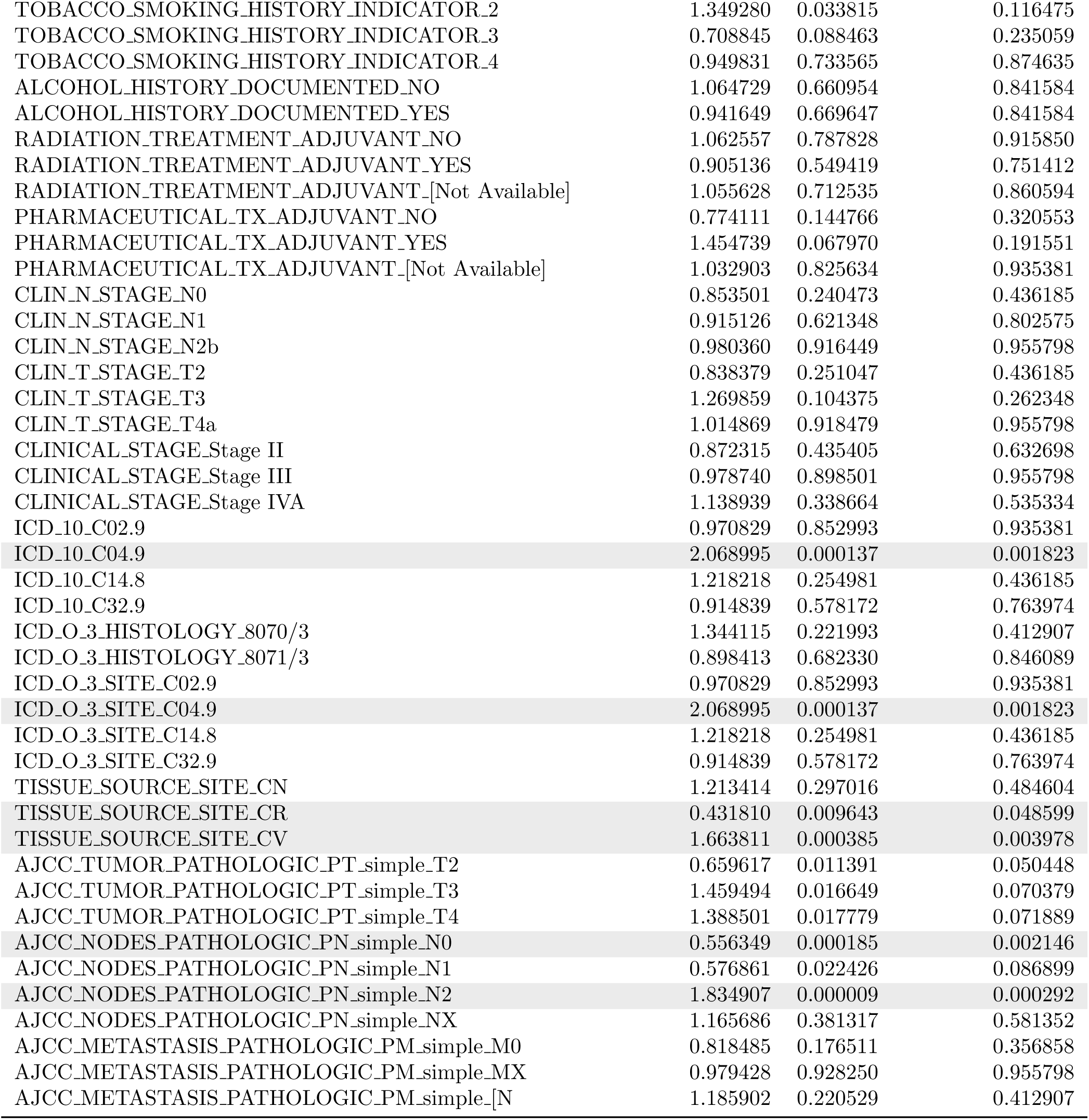
Clinical variables significance for single-variable cox model for HNSC (526 patients). Variable significantly associated with survival are shaded.

| Variable | Hazard ratio | P-value | Corrected P-value |
| --- | --- | --- | --- |
| INITIAL_PATHOLOGIC_DX_YEAR | 0.944473 | 0.000002 | 0.000085 |
| age_at_diagnosis | 1.019742 | 0.001594 | 0.012355 |
| PRIMARY_SITE_Floor of mouth | 1.633189 | 0.009866 | 0.048599 |
| PRIMARY_SITE_Larynx | 0.870495 | 0.392949 | 0.589424 |
| PRIMARY_SITE_Oral Cavity | 1.216594 | 0.257959 | 0.436185 |
| PRIMARY_SITE_Oral Tongue | 1.027363 | 0.864976 | 0.935381 |
| LATERALITY_Left | 0.997223 | 0.985683 | 0.985683 |
| LATERALITY_Right | 0.596169 | 0.001229 | 0.010391 |
| LATERALITY_[Not Available] | 1.487693 | 0.003515 | 0.021791 |
| PROSPECTIVE_COLLECTION_NO | 1.275968 | 0.136430 | 0.309463 |
| PROSPECTIVE_COLLECTION_YES | 0.796014 | 0.163125 | 0.337124 |
| RETROSPECTIVE_COLLECTION_NO | 0.796014 | 0.163125 | 0.337124 |
| RETROSPECTIVE_COLLECTION_YES | 1.275968 | 0.136430 | 0.309463 |
| SEX_Female | 1.382324 | 0.024394 | 0.087255 |
| SEX_Male | 0.723420 | 0.024394 | 0.087255 |
| RACE_WHITE | 0.742605 | 0.111804 | 0.273433 |
| ETHNICITY_NOT HISPANIC OR LATINO | 0.853619 | 0.414453 | 0.611812 |
| LYMPH_NODE_NECK_DISSECTION_INDICATOR_NO | 1.236683 | 0.203896 | 0.403454 |
| LYMPH_NODE_NECK_DISSECTION_INDICATOR_YES | 0.814286 | 0.215391 | 0.412907 |
| LYMPH_NODE_DISSECTION_METHOD_Functional (Limite... | 1.086460 | 0.540168 | 0.749786 |
| LYMPH_NODE_DISSECTION_METHOD_Modified Radical N... | 0.741422 | 0.093637 | 0.241895 |
| LYMPH_NODE_DISSECTION_METHOD_[Not Available] | 1.090921 | 0.583249 | 0.763974 |
| LYMPH_NODES_EXAMINED_NO | 1.470161 | 0.061855 | 0.179766 |
| LYMPH_NODES_EXAMINED_YES | 0.837805 | 0.290092 | 0.481761 |
| PATH_MARGIN_Negative | 0.777605 | 0.079280 | 0.216854 |
| PATH_MARGIN_Positive | 1.727697 | 0.002205 | 0.014645 |
| PATH_MARGIN_[Not Available] | 0.699698 | 0.158253 | 0.337124 |
| AJCC_STAGING_EDITION_6th | 1.010909 | 0.944167 | 0.955798 |
| AJCC_STAGING_EDITION_7th | 0.973421 | 0.855035 | 0.935381 |
| AJCC_PATHOLOGIC_TUMOR_STAGE_Stage II | 0.668524 | 0.054444 | 0.168777 |
| AJCC_PATHOLOGIC_TUMOR_STAGE_Stage III | 0.726051 | 0.114665 | 0.273433 |
| AJCC_PATHOLOGIC_TUMOR_STAGE_Stage IVA | 1.532143 | 0.001807 | 0.012924 |
| AJCC_PATHOLOGIC_TUMOR_STAGE_[Not Available] | 0.962185 | 0.849093 | 0.935381 |
| EXTRACAPSULAR_SPREAD_PATHOLOGIC_Microscopic<br>Ext... | 2.254494 | 0.000001 | 0.000085 |
| EXTRACAPSULAR_SPREAD_PATHOLOGIC_No Extranodal<br>E... | 0.548915 | 0.000017 | 0.000392 |
| EXTRACAPSULAR_SPREAD_PATHOLOGIC_[Not Available] | 0.990168 | 0.945521 | 0.955798 |
| GRADE_G1 | 0.657592 | 0.059833 | 0.179498 |
| GRADE_G2 | 1.431246 | 0.010999 | 0.050448 |
| GRADE_G3 | 0.957962 | 0.783689 | 0.915850 |
| ANGIOLYMPHATIC_INVASION_NO | 0.693371 | 0.009929 | 0.048599 |
| ANGIOLYMPHATIC_INVASION_YES | 1.484568 | 0.008174 | 0.047509 |
| ANGIOLYMPHATIC_INVASION_[Not Available] | 1.057948 | 0.692475 | 0.847371 |
| PERINEURAL_INVASION_NO | 0.516015 | 0.000030 | 0.000565 |
| PERINEURAL_INVASION_YES | 1.588730 | 0.000670 | 0.006230 |
| PERINEURAL_INVASION_[Not Available] | 1.147166 | 0.339620 | 0.535334 |
| HPV_STATUS_P16_Negative | 0.856744 | 0.477818 | 0.673289 |
| HPV_STATUS_P16_[Not Available] | 1.469555 | 0.052450 | 0.168202 |
| HPV_STATUS_ISH_Negative | 0.842503 | 0.448772 | 0.642089 |
| HPV_STATUS_ISH_[Not Available] | 1.564703 | 0.043660 | 0.145013 |
| TOBACCO_SMOKING_HISTORY_INDICATOR_1 | 0.853226 | 0.347479 | 0.538592 |
| TOBACCO_SMOKING_HISTORY_INDICATOR_2 | 1.349280 | 0.033815 | 0.116475 |
| TOBACCO_SMOKING_HISTORY_INDICATOR_3 | 0.708845 | 0.088463 | 0.235059 |
| TOBACCO_SMOKING_HISTORY_INDICATOR_4 | 0.949831 | 0.733565 | 0.874635 |
| ALCOHOL_HISTORY_DOCUMENTED_NO | 1.064729 | 0.660954 | 0.841584 |
| ALCOHOL_HISTORY_DOCUMENTED_YES | 0.941649 | 0.669647 | 0.841584 |
| RADIATION_TREATMENT_ADJUVANT_NO | 1.062557 | 0.787828 | 0.915850 |
| RADIATION_TREATMENT_ADJUVANT_YES | 0.905136 | 0.549419 | 0.751412 |
| RADIATION_TREATMENT_ADJUVANT_[Not Available] | 1.055628 | 0.712535 | 0.860594 |
| PHARMACEUTICAL_TX_ADJUVANT_NO | 0.774111 | 0.144766 | 0.320553 |
| PHARMACEUTICAL_TX_ADJUVANT_YES | 1.454739 | 0.067970 | 0.191551 |
| PHARMACEUTICAL_TX_ADJUVANT_[Not Available] | 1.032903 | 0.825634 | 0.935381 |
| CLIN_N_STAGE_N0 | 0.853501 | 0.240473 | 0.436185 |
| CLIN_N_STAGE_N1 | 0.915126 | 0.621348 | 0.802575 |
| CLIN_N_STAGE_N2b | 0.980360 | 0.916449 | 0.955798 |
| CLIN_T_STAGE_T2 | 0.838379 | 0.251047 | 0.436185 |
| CLIN_T_STAGE_T3 | 1.269859 | 0.104375 | 0.262348 |
| CLIN_T_STAGE_T4a | 1.014869 | 0.918479 | 0.955798 |
| CLINICAL_STAGE_Stage II | 0.872315 | 0.435405 | 0.632698 |
| CLINICAL_STAGE_Stage III | 0.978740 | 0.898501 | 0.955798 |
| CLINICAL_STAGE_Stage IVA | 1.138939 | 0.338664 | 0.535334 |
| ICD_10_C02.9 | 0.970829 | 0.852993 | 0.935381 |
| ICD_10_C04.9 | 2.068995 | 0.000137 | 0.001823 |
| ICD_10_C14.8 | 1.218218 | 0.254981 | 0.436185 |
| ICD_10_C32.9 | 0.914839 | 0.578172 | 0.763974 |
| ICD_O_3_HISTOLOGY_8070/3 | 1.344115 | 0.221993 | 0.412907 |
| ICD_O_3_HISTOLOGY_8071/3 | 0.898413 | 0.682330 | 0.846089 |
| ICD_O_3_SITE_C02.9 | 0.970829 | 0.852993 | 0.935381 |
| ICD_O_3_SITE_C04.9 | 2.068995 | 0.000137 | 0.001823 |
| ICD_O_3_SITE_C14.8 | 1.218218 | 0.254981 | 0.436185 |
| ICD_O_3_SITE_C32.9 | 0.914839 | 0.578172 | 0.763974 |
| TISSUE_SOURCE_SITE_CN | 1.213414 | 0.297016 | 0.484604 |
| TISSUE_SOURCE_SITE_CR | 0.431810 | 0.009643 | 0.048599 |
| TISSUE_SOURCE_SITE_CV | 1.663811 | 0.000385 | 0.003978 |
| AJCC_TUMOR_PATHOLOGIC_PT_simple_T2 | 0.659617 | 0.011391 | 0.050448 |
| AJCC_TUMOR_PATHOLOGIC_PT_simple_T3 | 1.459494 | 0.016649 | 0.070379 |
| AJCC_TUMOR_PATHOLOGIC_PT_simple_T4 | 1.388501 | 0.017779 | 0.071889 |
| AJCC_NODES_PATHOLOGIC_PN_simple_N0 | 0.556349 | 0.000185 | 0.002146 |
| AJCC_NODES_PATHOLOGIC_PN_simple_N1 | 0.576861 | 0.022426 | 0.086899 |
| AJCC_NODES_PATHOLOGIC_PN_simple_N2 | 1.834907 | 0.000009 | 0.000292 |
| AJCC_NODES_PATHOLOGIC_PN_simple_NX | 1.165686 | 0.381317 | 0.581352 |
| AJCC_METASTASIS_PATHOLOGIC_PM_simple_M0 | 0.818485 | 0.176511 | 0.356858 |
| AJCC_METASTASIS_PATHOLOGIC_PM_simple_MX | 0.979428 | 0.928250 | 0.955798 |
| AJCC_METASTASIS_PATHOLOGIC_PM_simple_[N | 1.185902 | 0.220529 | 0.412907 |

**Table S5:**
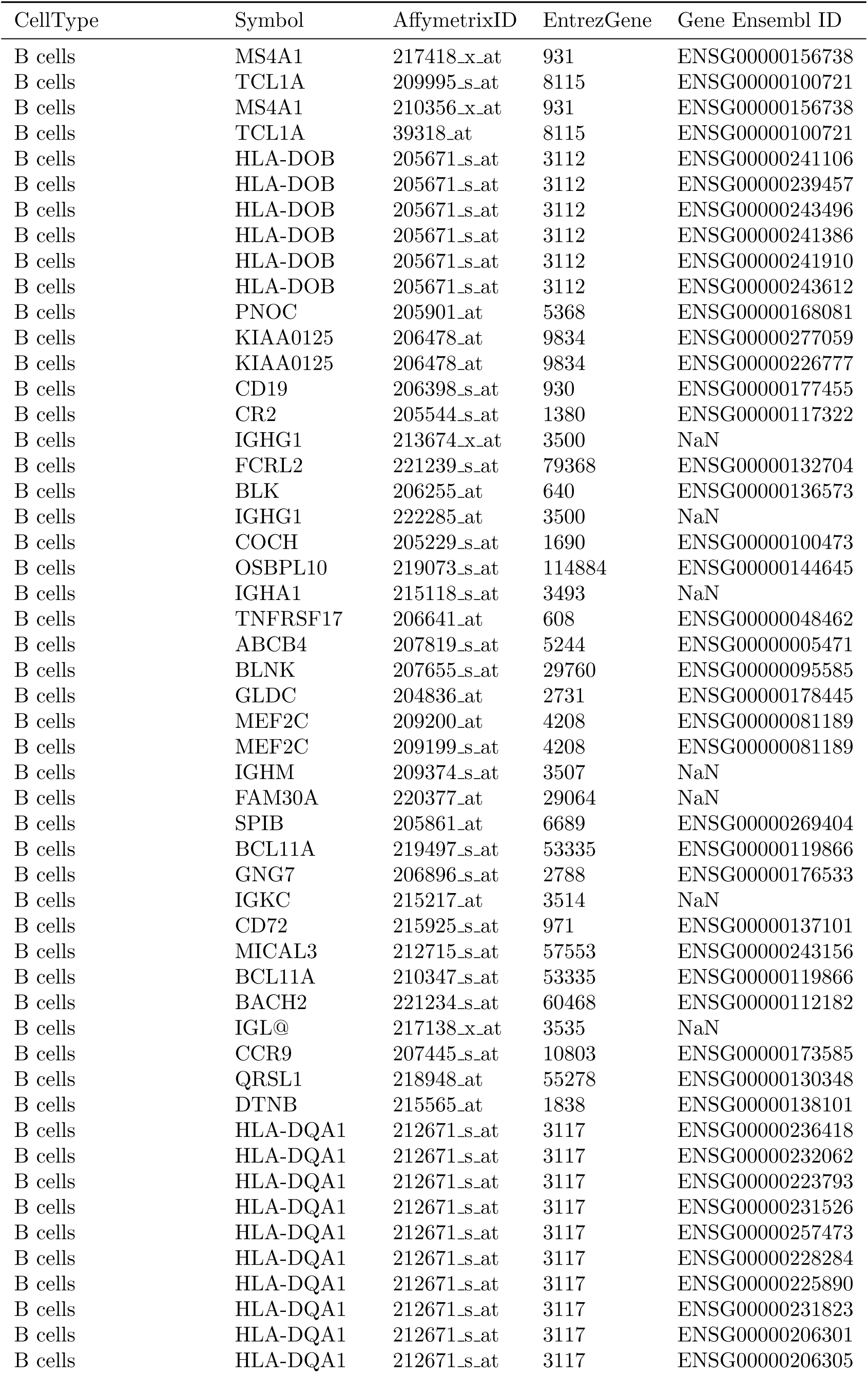
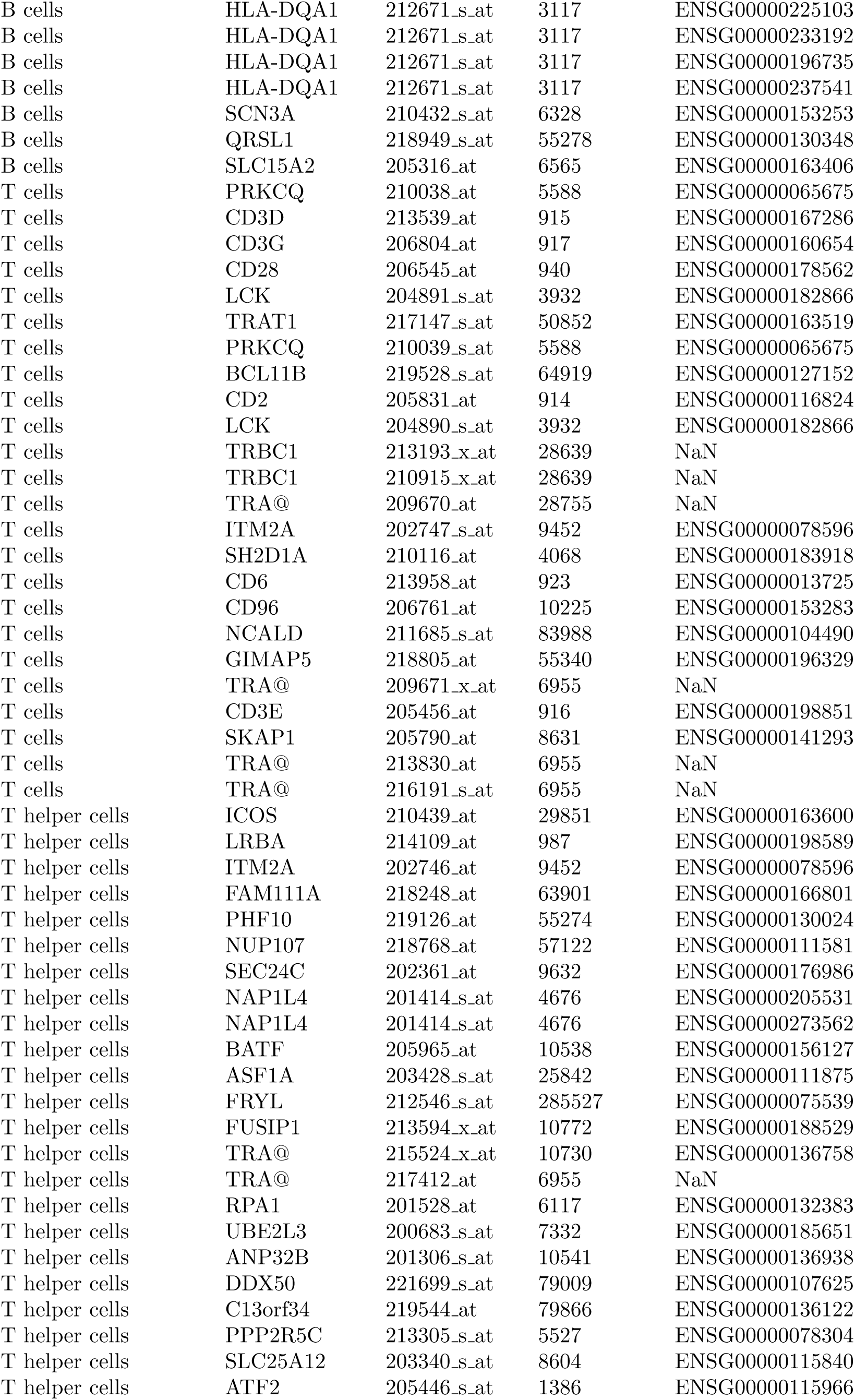
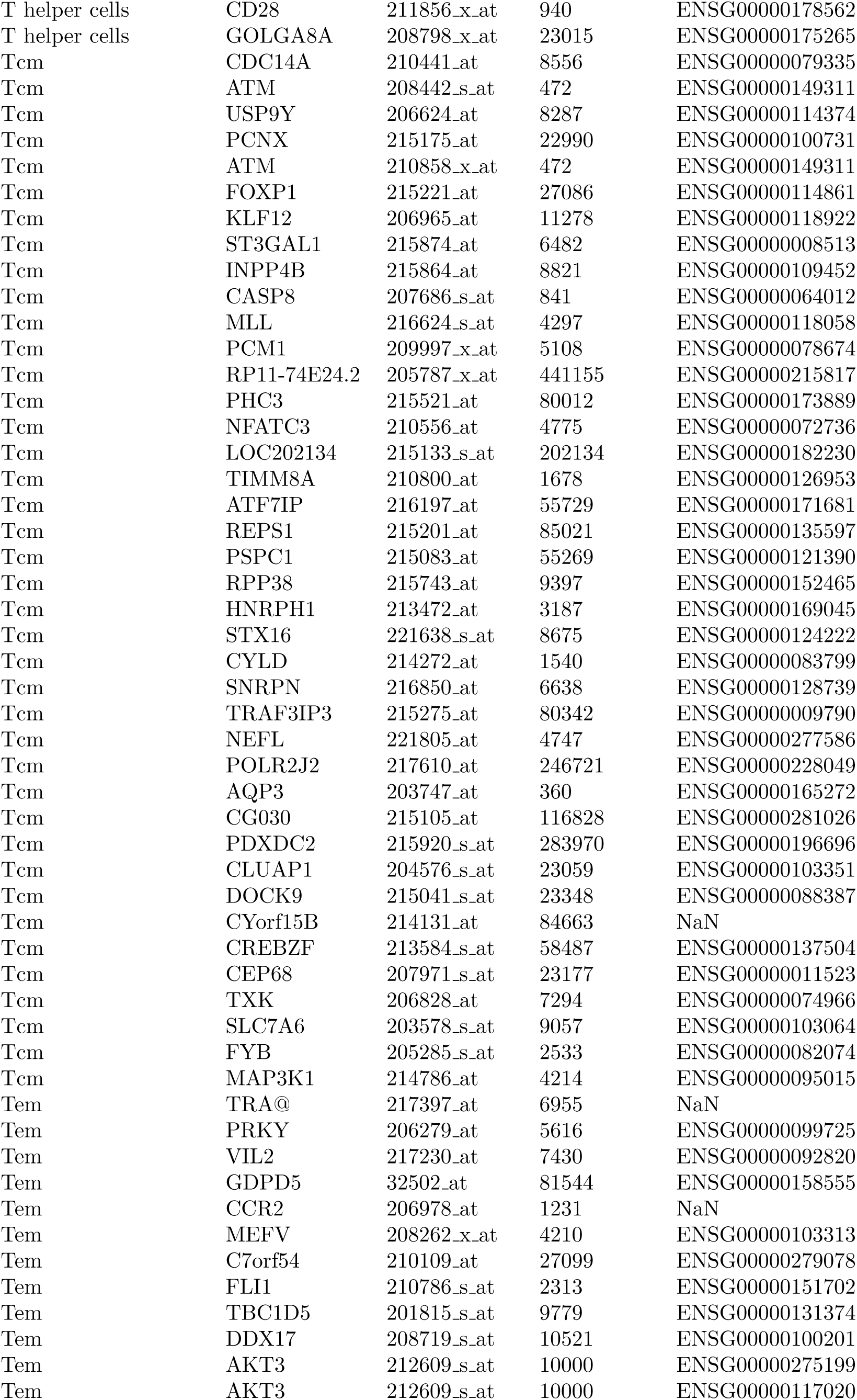
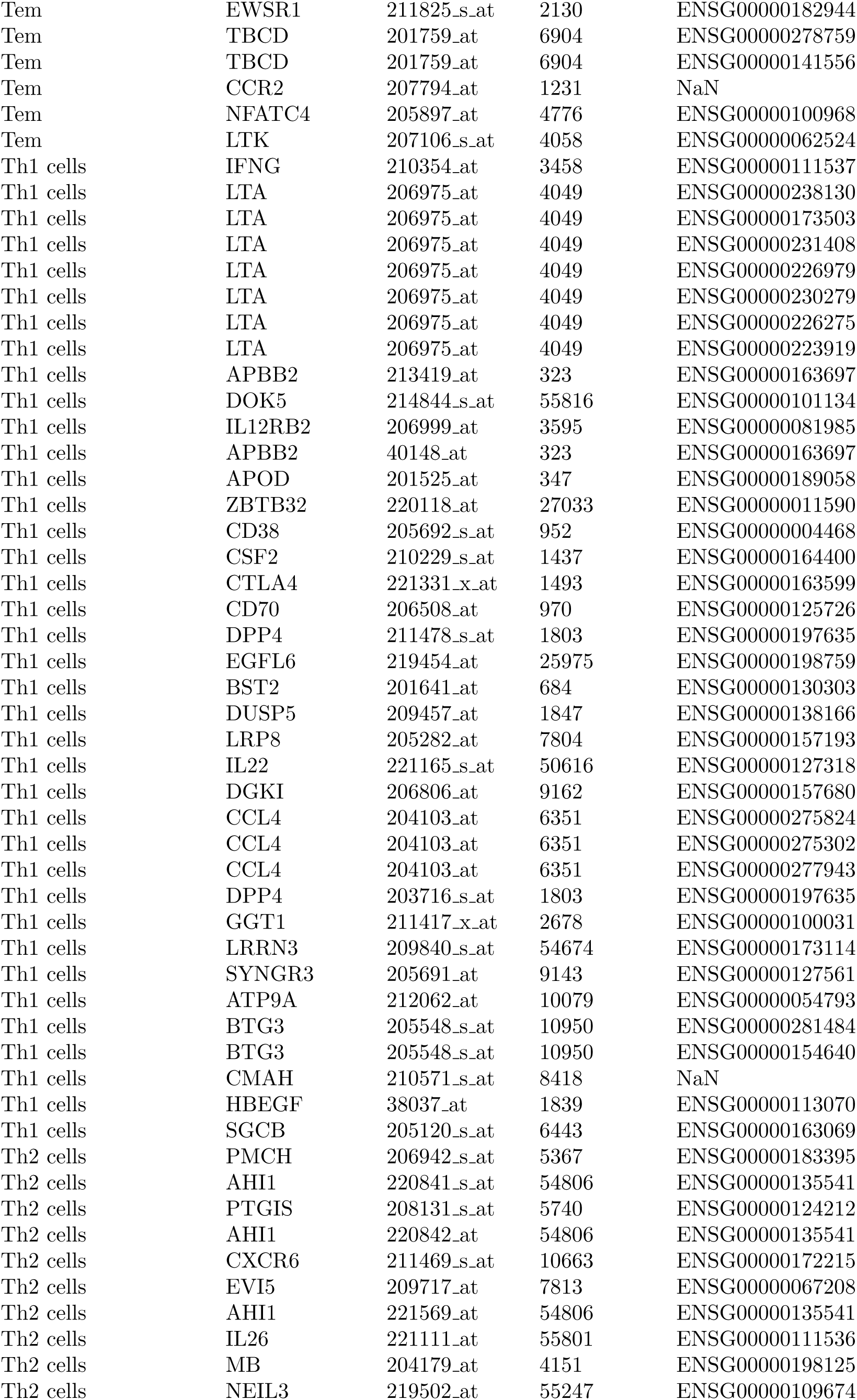
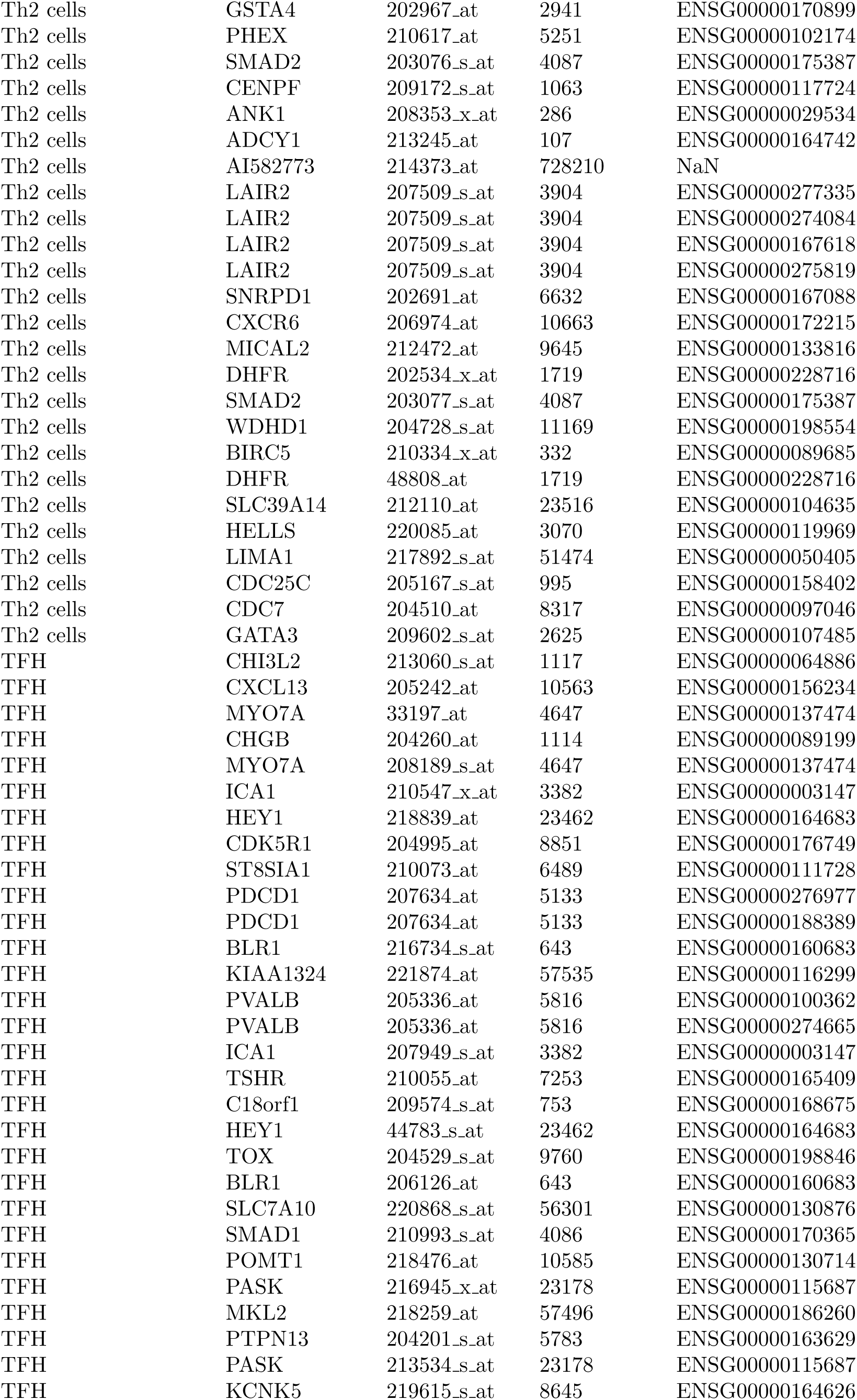
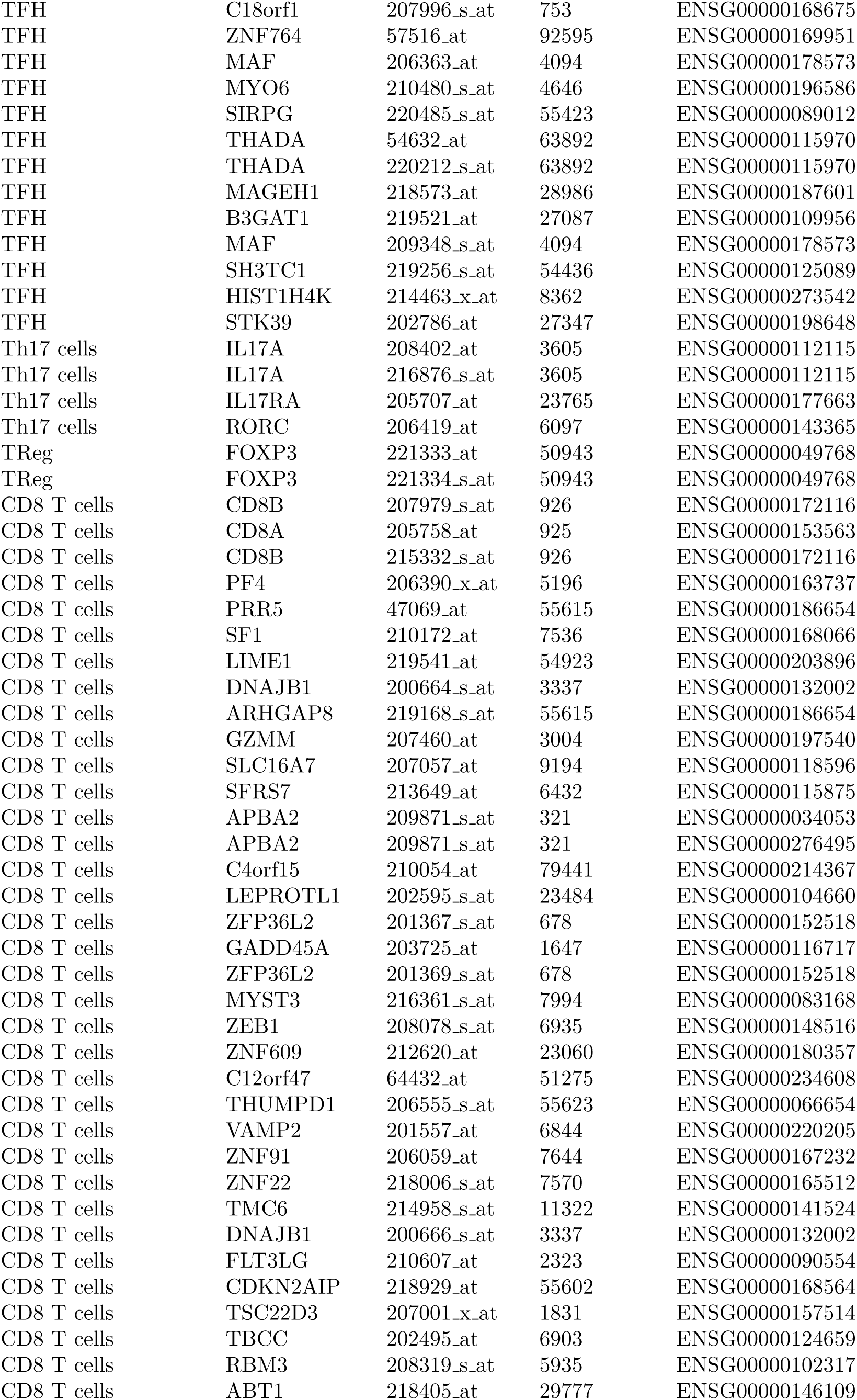
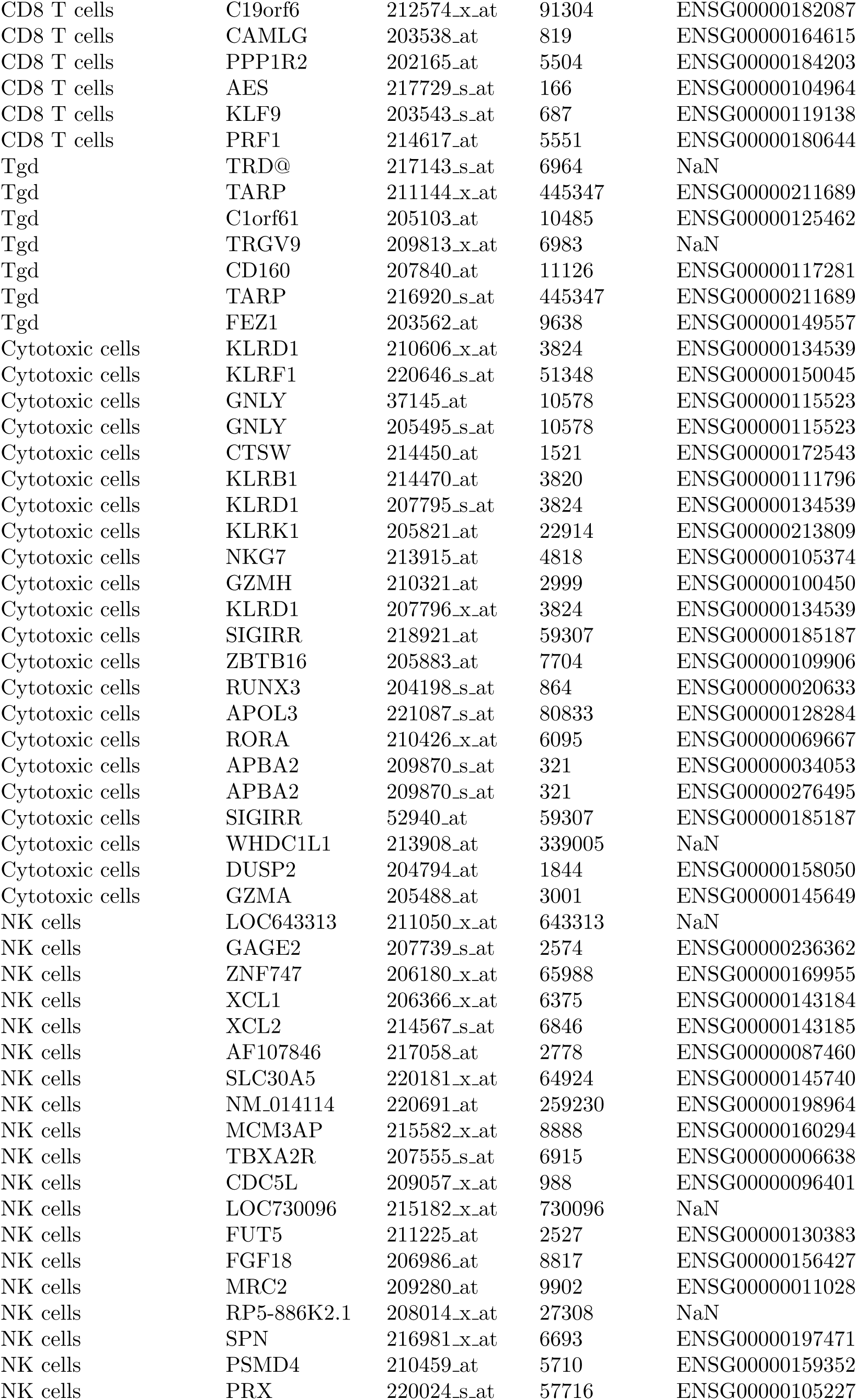
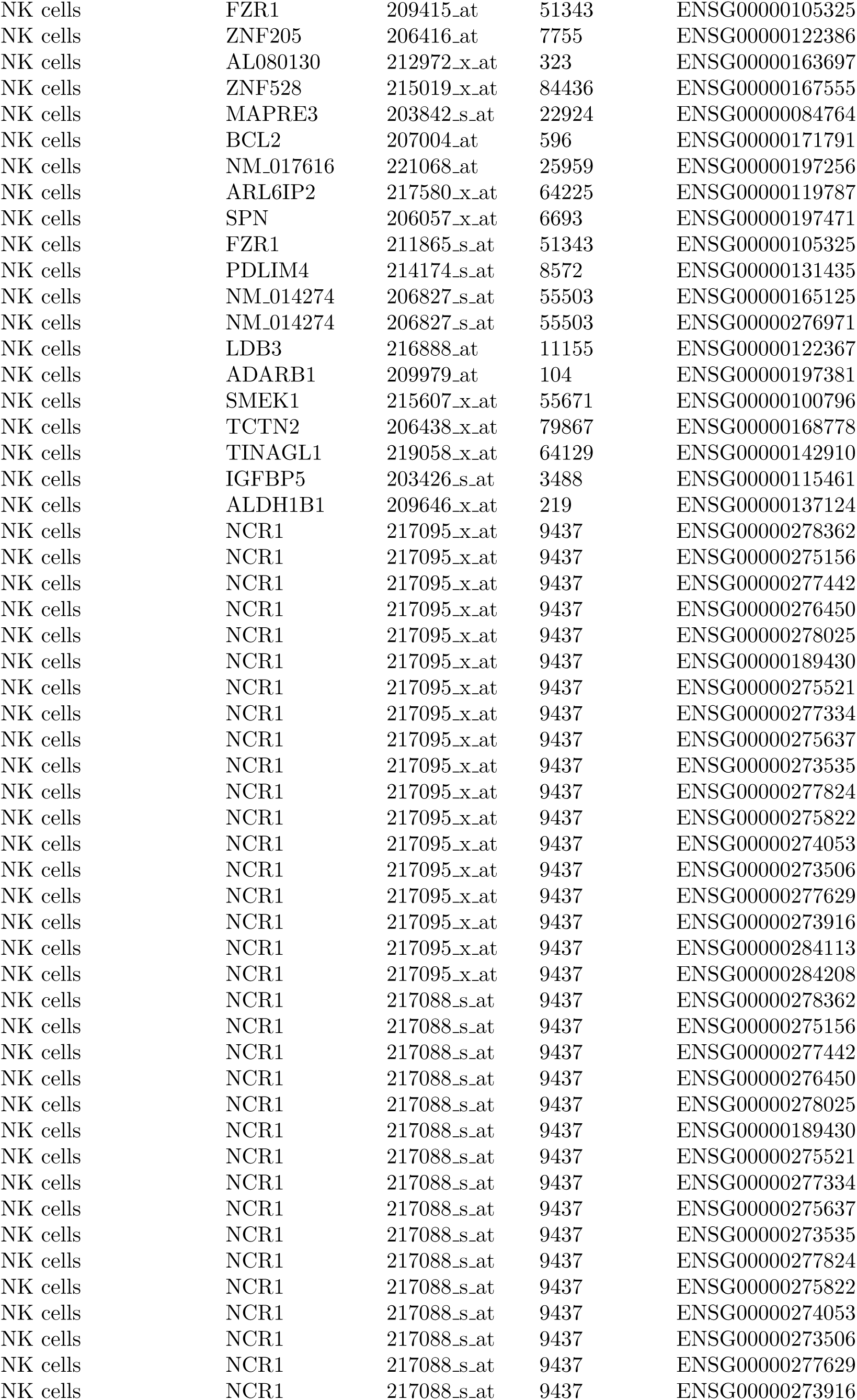
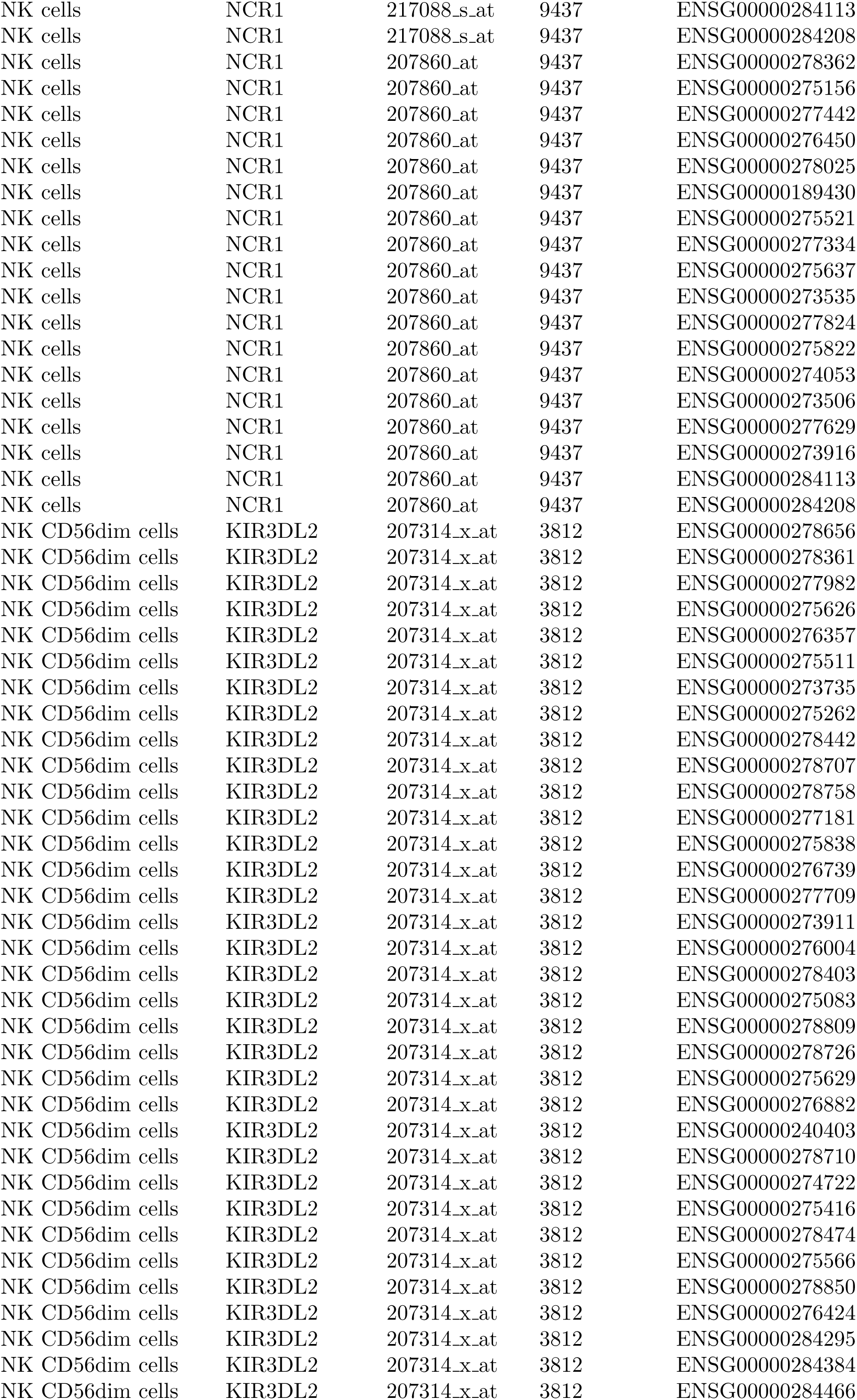
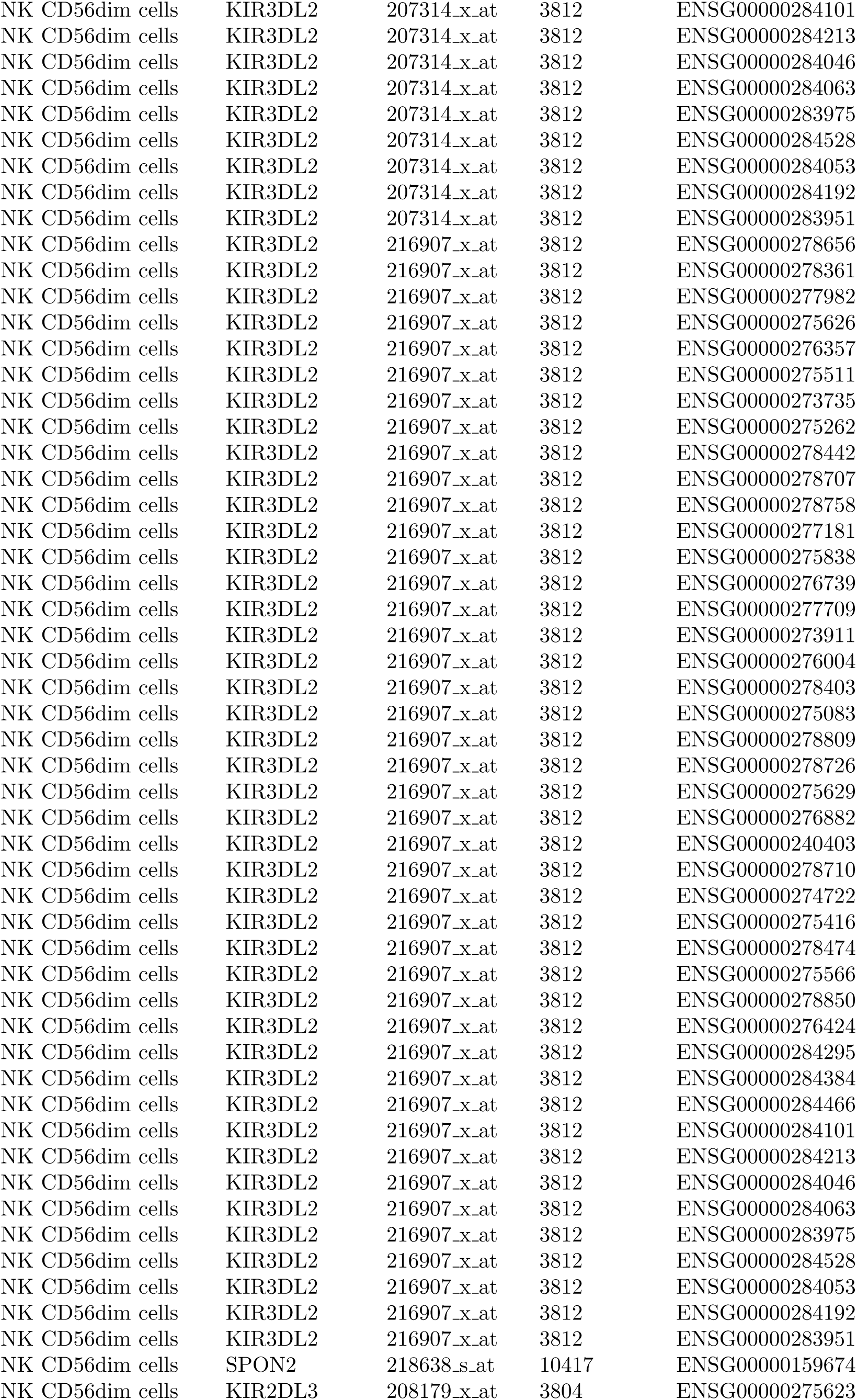
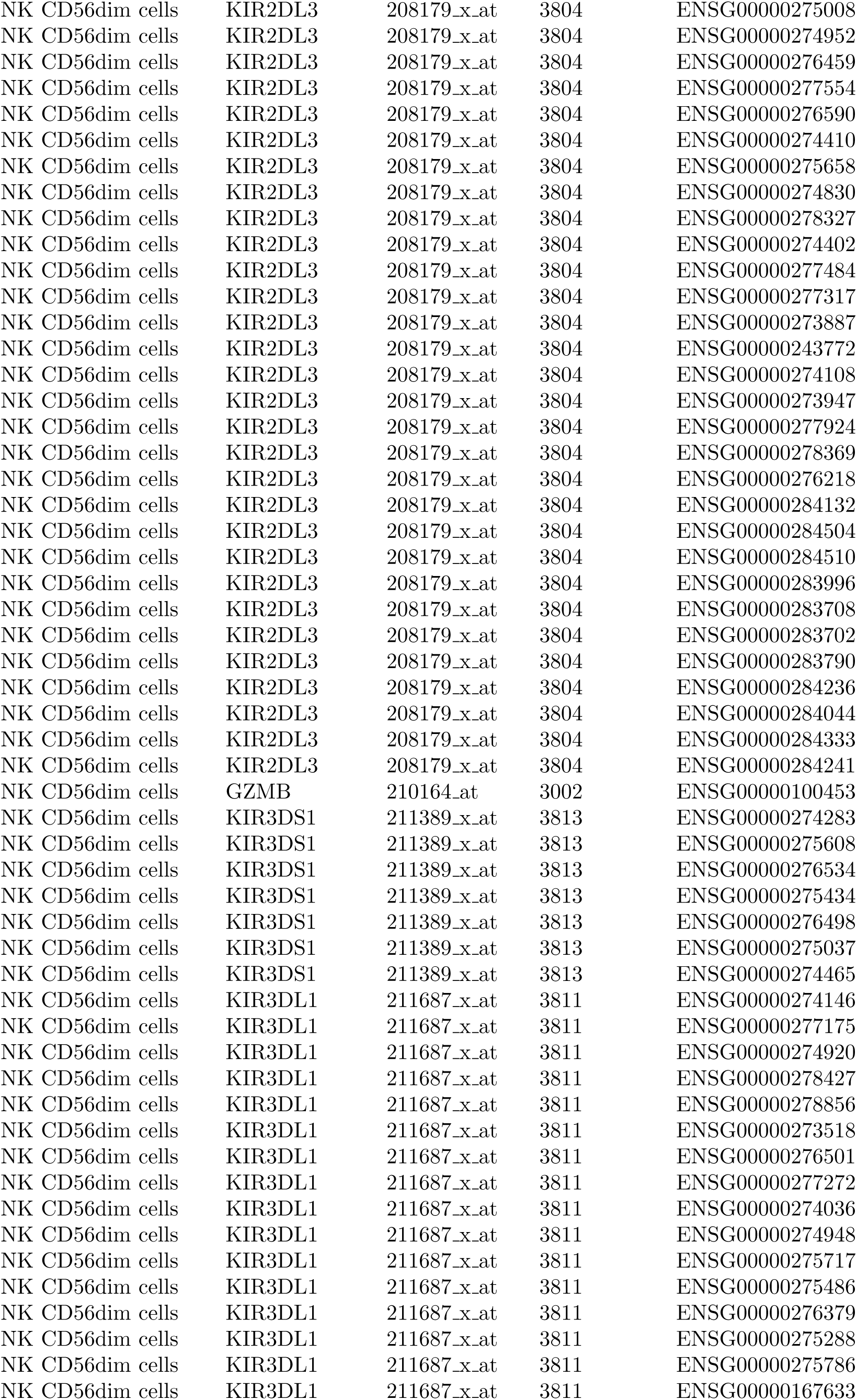
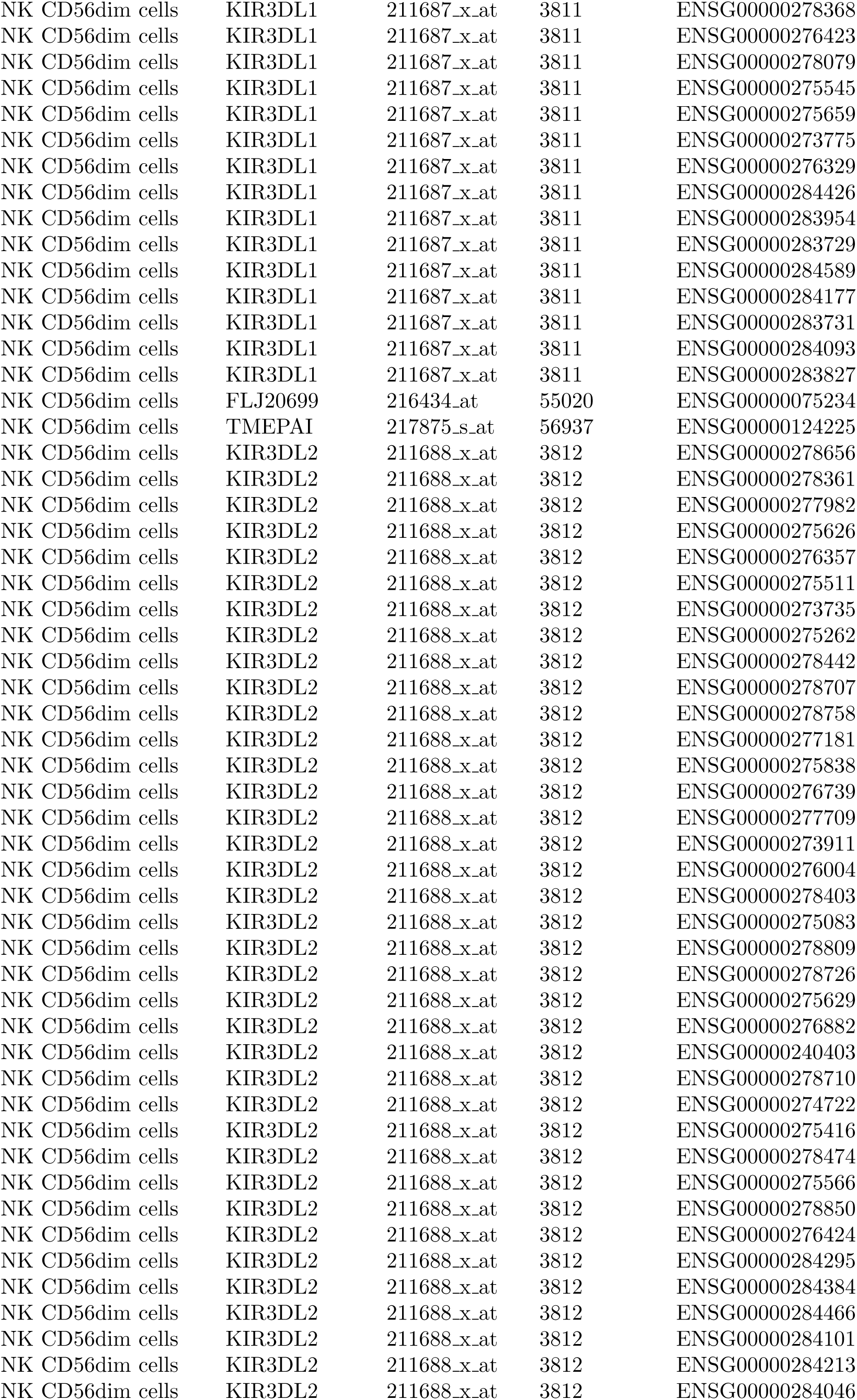
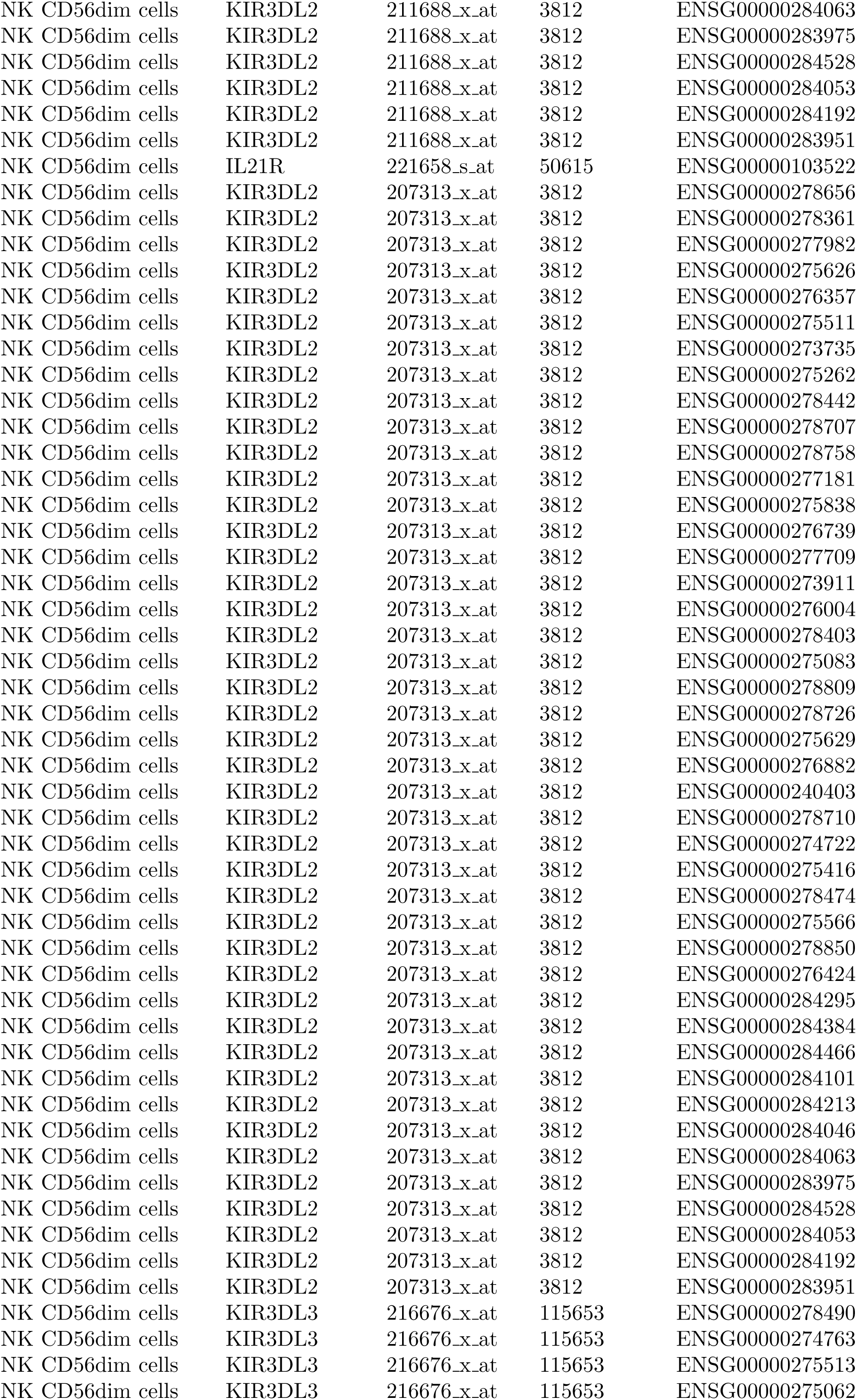
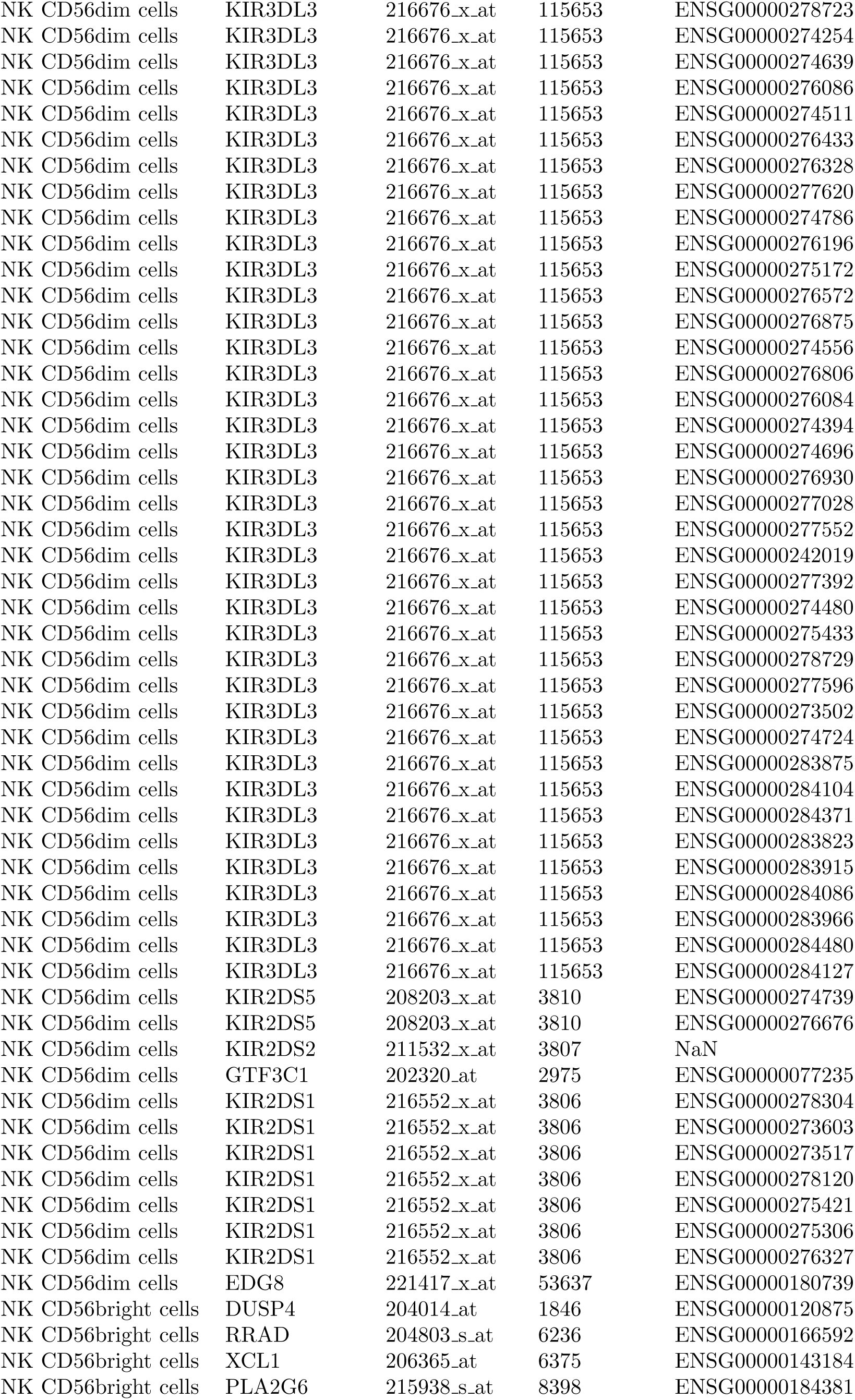
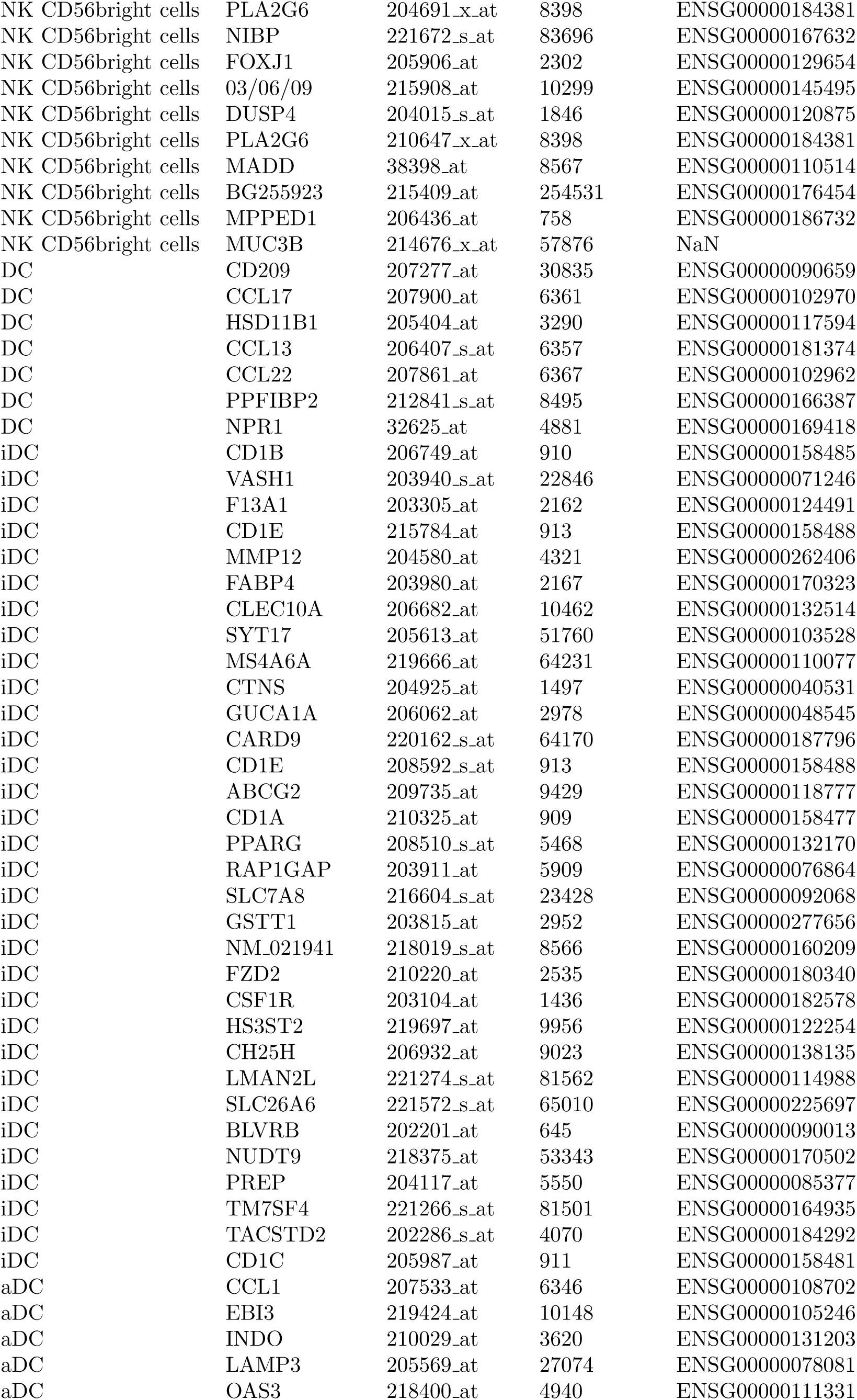
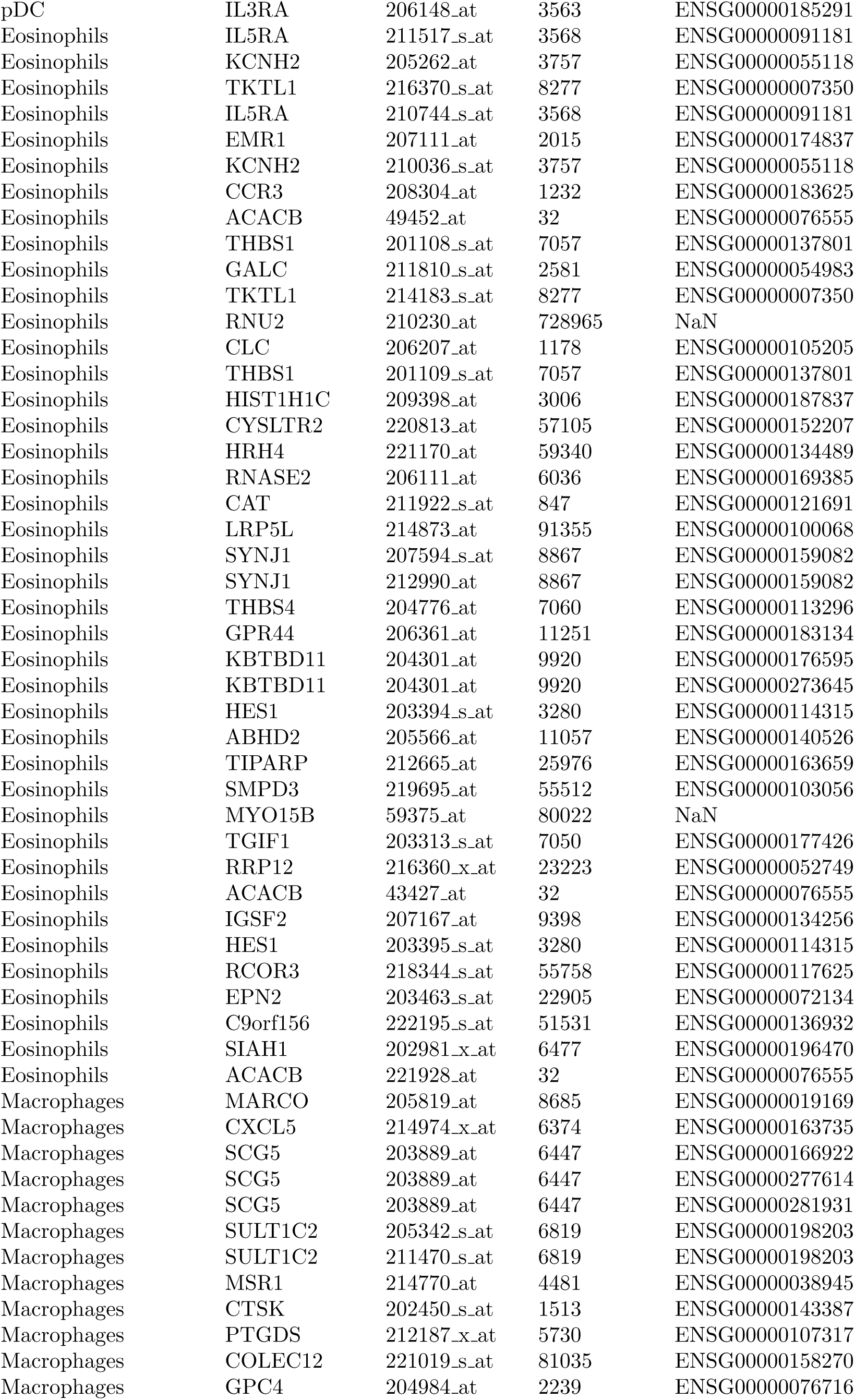
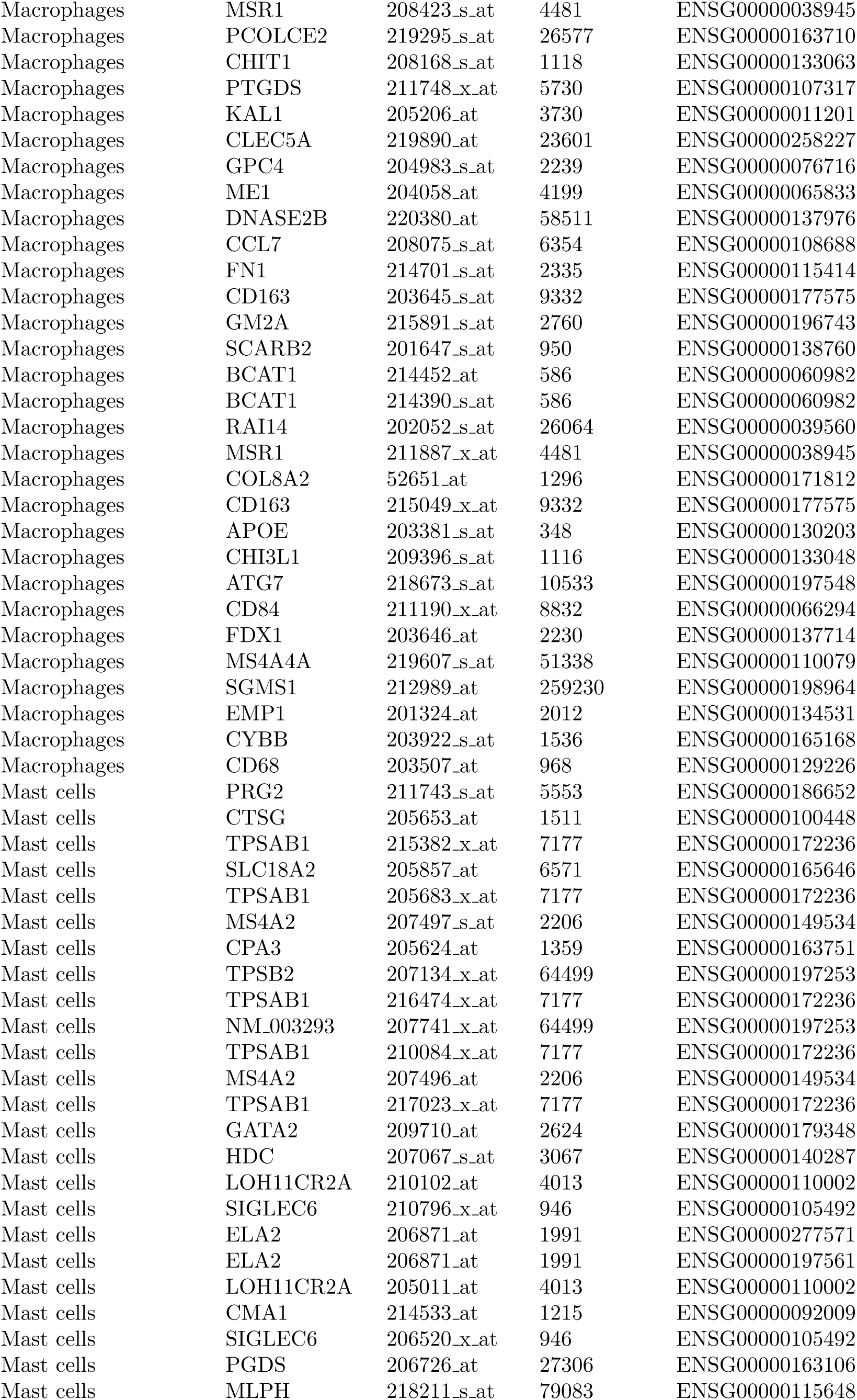
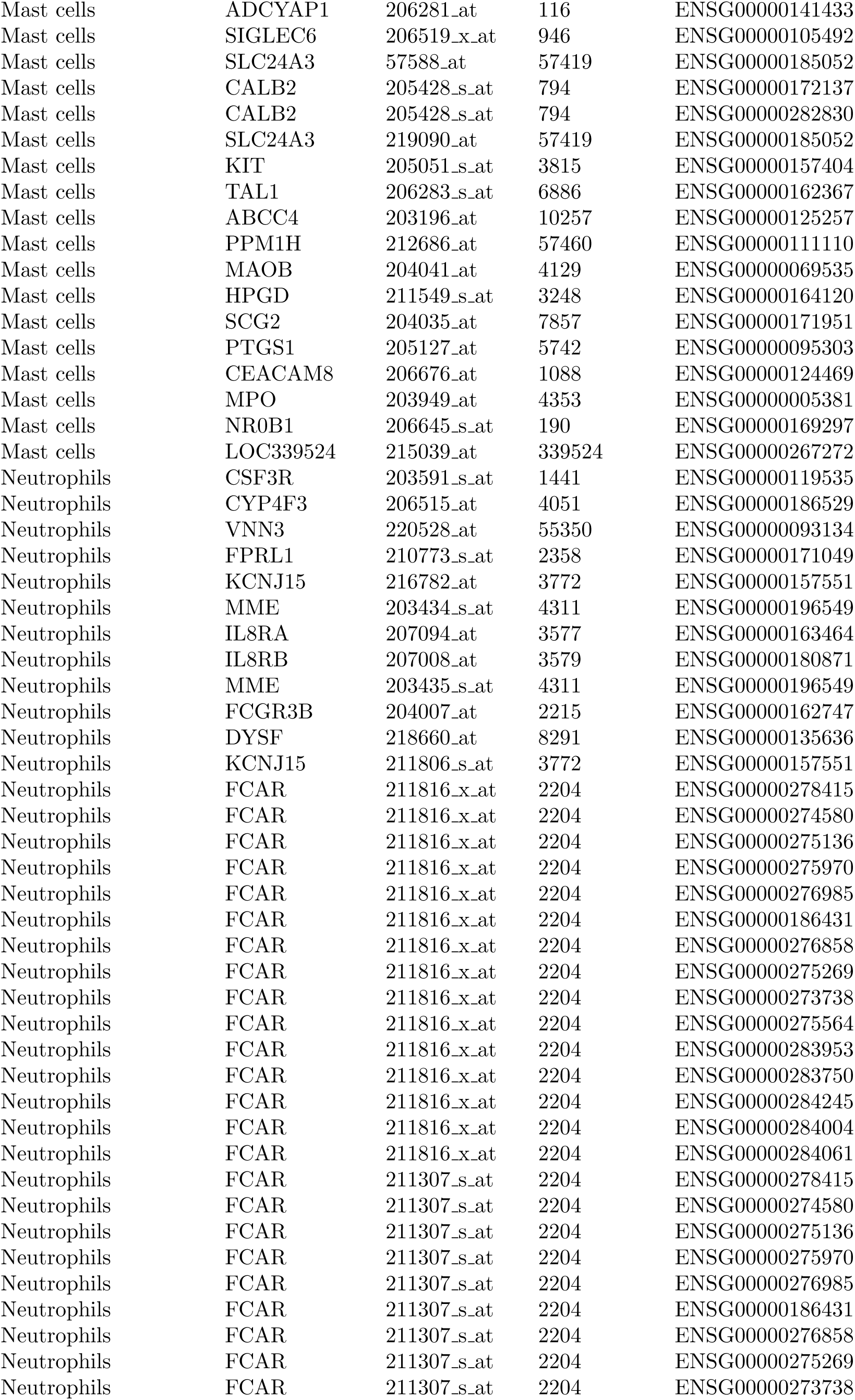
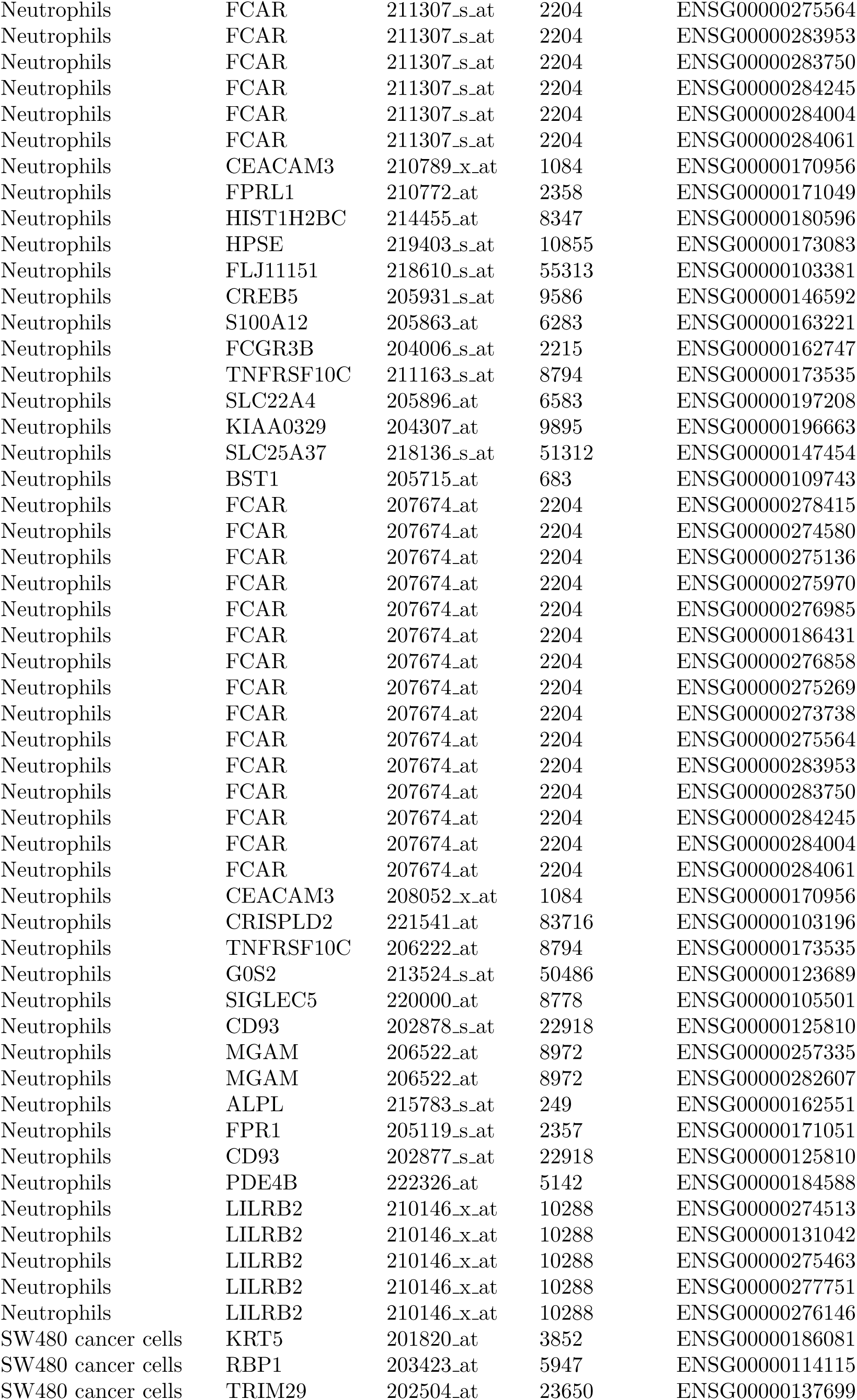
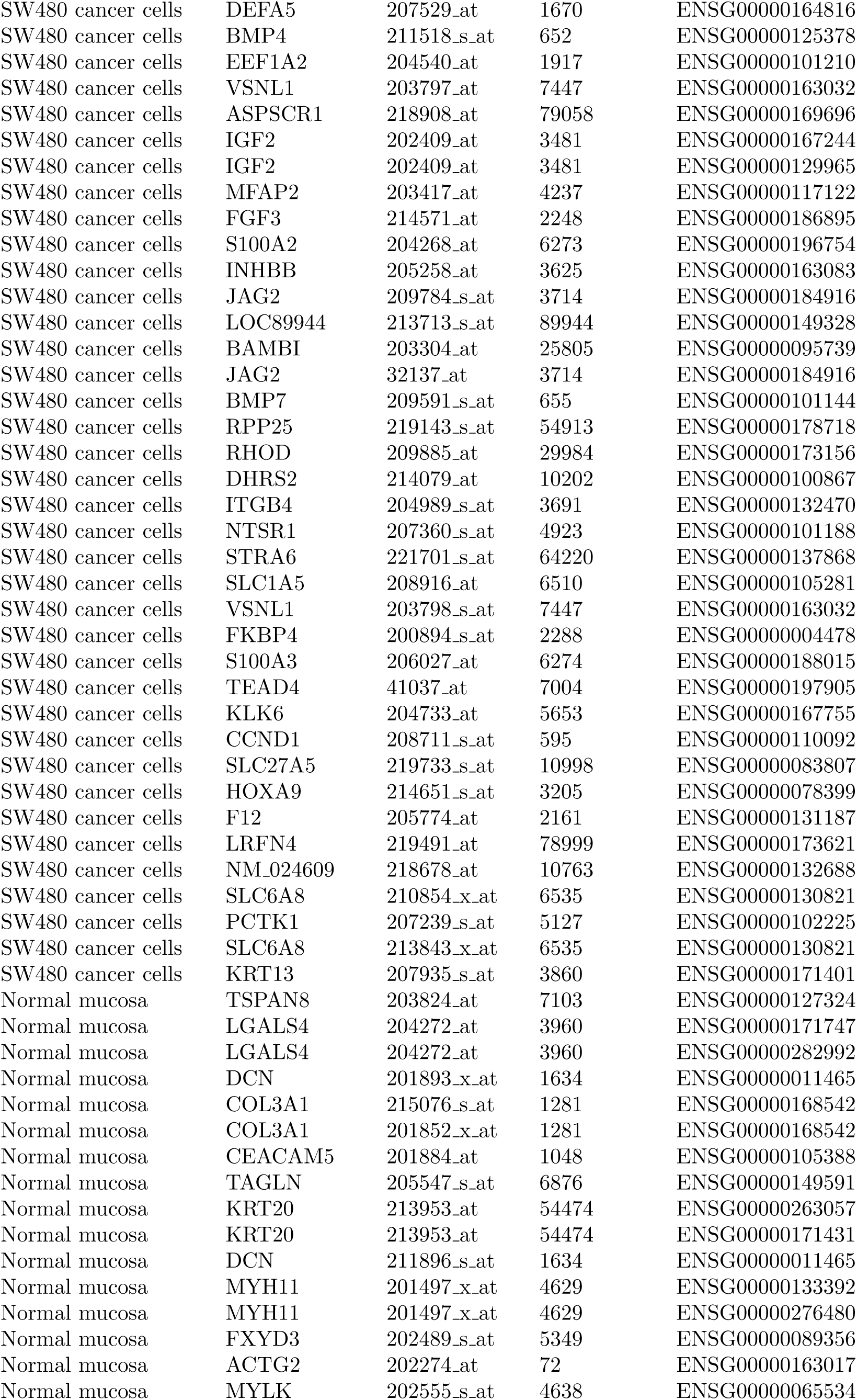
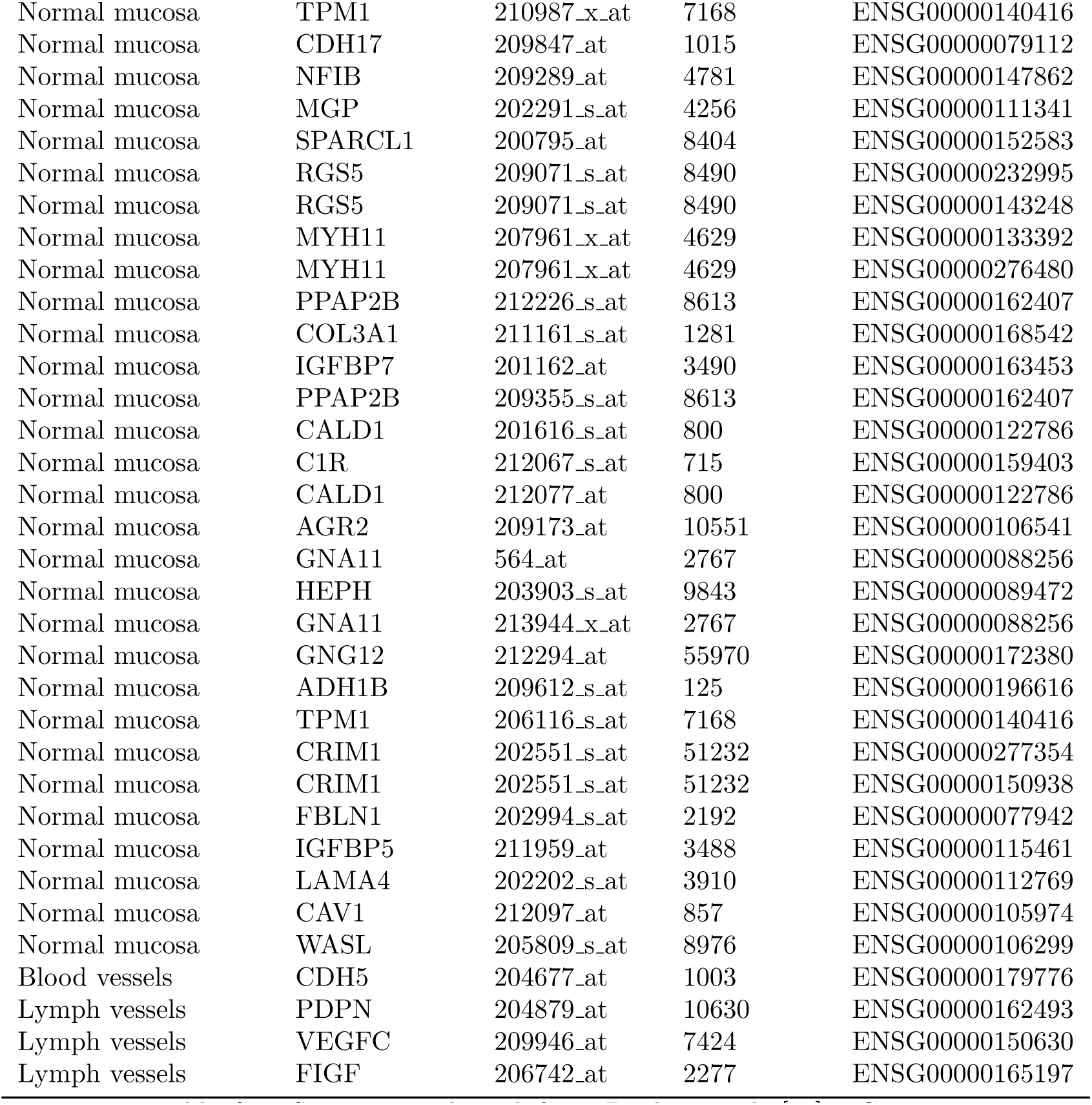
Signatures adapted from Bindea et al. [38]. Genes Id were matched from tables available at https://github.com/judithabk6/ITH_TCGA/tree/master/external_data

